# Elucidating the neuropathological and molecular heterogeneity of amyloid-β and tau in Alzheimer’s disease through machine learning and transcriptomic integration

**DOI:** 10.1101/2024.10.16.618708

**Authors:** Kanhao Zhao, Hua Xie, Tovia Jacobs, Naomi L. Gaggi, Juan Fortea, Nancy B. Carlisle, Gregory A. Fonzo, Kilian M. Pohl, Ricardo S. Osorio, Yu Zhang, the ADNI Study Group and the PREVENT-AD Research Group

## Abstract

Discerning functional brain network variations related to neuropathological aggregates in Alzheimer’s disease (AD), including amyloid-β (Aβ) and phosphorylated tau (p-tau), is crucial for understanding their link to cognitive decline and underlying molecular mechanisms. However, these variations are often confounded by normal aging-related changes, complicating interpretation. To address this challenge, we first defined Alzheimer’s continuum cases (Aβ positive (A^+^), n = 129) and normal elderly (Aβ negative (A^-^), n = 160) using cerebral spinal fluid amyloid levels, and then applied a novel deep learning approach to resting-state connectivity using functional magnetic resonance imaging (fMRI) of the 289 subjects to disentangle A^+^-specific dimensions in brain network alterations from those shared with A^-^ individuals. The identified A^+^-specific dimensions were further refined to predict individual Aβ and p-tau levels separately. We observed that resulting brain signatures, defined from A^+^-specific dimensions for predicting these two CSF biomarkers, were both attributed to the right superior temporal and anterior cingulate cortices and associated with attention and memory domains. When linking the brain signatures to gene expression data from a public transcriptomic atlas, we found that the brain signatures were associated with molecular pathways involving synaptic dysfunction and disruptions in pathways containing activity of excitatory neurons, astrocytes, and microglia. For A^-^-shared dimensions, the Aβ-linked brain signature involved the left fusiform and right middle cingulate cortices, correlating with the language cognitive measurement and language-related molecular pathways. The p-tau-linked signature predominantly involved the right insula and inferior temporal cortices, correlating with the aging-related molecular pathways. Collectively, our findings provided new insights in understanding of Alzheimer’s continuum pathological biomarkers.

## Introduction

Despite millions of individuals and their caregivers being affected by Alzheimer’s disease (AD)^1^ worldwide, and with its prevalence expected to grow as the population ages, our understanding of the disease’s natural history and progression remains limited^2^. This lack of knowledge presents a significant challenge in developing preventative strategies, as it hinders our ability to adequately predict the course of the disease and implement effective interventions.

Amyloid-beta (Aβ) plaques and neurofibrillary tangles (aggregates of hyperphosphorylated tau (p-tau)) are key neuropathological factors in the development of AD^3^. The cascading effects of Aβ deposition and p-tau propagation lead to a sequence of detrimental outcomes, including neuronal loss, disruption of brain networks, neuroinflammation, vascular damage, and ultimately, dementia^4^. Specifically, the disruptions in cognition-associated brain networks in AD have been highlighted by studies of functional connectivity extracted from functional magnetic resonance imaging (fMRI)^5,6^. However, AD-specific brain functional abnormalities, caused by Aβ and tau accumulations^7^ can be obscured by those observed in normal aging, as brain abnormalities in AD may result from a combination of biological processes involving both normal aging and AD-specific pathology^8–10^. This challenge hinders our understanding of AD-specific pathological processes and how they contribute to these functional disruptions and complicates the development of precise therapeutic strategies^11^. Consequently, there is an urgent need for the development of innovative analytical techniques to discern ‘AD-specific variations’ from those of normal aging.

A better understanding of AD-specific and aging-shared pathological variations would be beneficial to understanding genetic factors that regulate the expression of AD neuropathology^12,13^. The Allen Human Brain Atlas (AHBA)^14^, which contains messenger RNA expression information across various brain regions, made the gene analysis for AD feasible. This atlas has become a crucial tool for investigating the association between gene expression profiles and brain attributes, such as cortical thickness, and fractional anisotropy in schizophrenia, autism, and major depressive disorder^15–19^. A recent study used AHBA for normal elderly to explore the role of genetic traits in tau propagation progression and revealed that the apolipoprotein E (*APOE*) and glutamatergic synaptic genes (*SLC1A2*) were critical in determining tau spreading^20^. However, applying this atlas to AD research, specifically to investigate how regional differences in gene expression within the brain contribute to the variations in functional connectivity associated with AD neuropathology, remains largely unexplored.

To disentangle brain functional variations specific to AD, we propose a cutting-edge deep learning framework in conjunction with resting-state fMRI connectivity (Figure S1). Our framework utilized a contrastive learning technique^21^, which differentiates between target and background datasets to separate entangled variations into different dimensions. This technique has been successfully applied to segregate autism-specific neuroanatomical and functional connectivity features from those shared with typically developing individuals^22,23^. Given the importance of topological abnormalities in AD pathology^24^, we represented functional brain networks as graphs and embedded graph convolutional networks (GCN) within the proposed contrastive learning setup^25,26^, forming our contrastive GCN (cGCN) model. We applied the model to data from two cohorts (n = 289): the Alzheimer’s Disease Neuroimaging Initiative (ADNI, n = 161)^27^ and the Pre-symptomatic Evaluation of Experimental or Novel Treatments for AD (PREVENT-AD, n = 128)^28^. We classified participants based on their CSF Aβ levels into either Alzheimer’s continuum cases (Aβ positive, A^+^) or controls (Aβ negative, A^-^) and trained our model to disentangle A^+^-specific dimensions in brain network alterations from those shared with A^-^ subjects. Compared to categorizations based on cognitive impairment (e.g., normal vs. dementia) or tau burden, the Aβ status-based contrastive framework enables a better screening of the AD biological continuum since the Aβ pathway is thought to represent one of the earliest pathophysiological events in AD^29,30^ and fMRI is a suitable biomarker of Alzheimer’s continuum^31,32^. Additionally, including A^-^ subjects with mild cognitive impairment (e.g., A^-^ MCI patients) may help differentiate between cognitive decline due to Aβ pathology and cognitive decline resulting from other factors, such as vascular contributions, Lewy body pathology, or depression, thus better accounting for clinical heterogeneity. The uncovered dimensions from our model were further refined to identify brain signatures that predict individual CSF Aβ_1-42_ and p-tau_181_ levels separately. Rigorous cross-validation and statistical analyses demonstrated the superior prediction accuracy of A^+^-specific dimensions identified by our model in comparison to conventional machine learning algorithms. To understand underlying molecular pathways of identified brain signatures, we applied partial least squares (PLS) regression, followed by gene set enrichment analysis (GSEA), to examine their associations with gene expression data from the ABHA. Finally, considering AD’s association with dysfunction of cell types including astrocytes, microglia, and oligodendrocytes^33,34^, we investigated the relationships between dysfunction of seven canonical neuron types and brain signatures using multi-gene-list meta-analysis.

## Results

### Brain signatures uncovered by cGCN for predicting neuropathological proteins

To train our cGCN model to obtain A^+^-specific features, we defined a target group (A^+^, n = 129) based on CSF Aβ_1-42_ < 976.6 pg/ml^35,36^ and a background group comprising A^-^ (n = 160) individuals based on CSF Aβ_1-42_ > 976.6 pg/ml (Table S1). We then constructed a brain graph based on z-scored functional connections for each individual (See Methods section for details). We contrasted the brain graph features between A^+^ and A^-^ groups to encode the features into A^+^-specific and A^-^-shared dimensions and demonstrate the ability of A^+^-specific dimension in predicting Aβ and p-tau pathology in the A^+^ group. The observed and predicted scores for Aβ and p-tau expression, evaluated using R^2^ and Pearson correlation (r) metrics, were significantly correlated across ten repetitions of 5-fold cross- validated models (Figures 1 (A, D); For Aβ expression, R^2^ = 0.19, r = 0.46, p < 0.0001; For p-tau expression, R^2^ = 0.26, r = 0.51, p < 0.0001).

**Figure 1.**
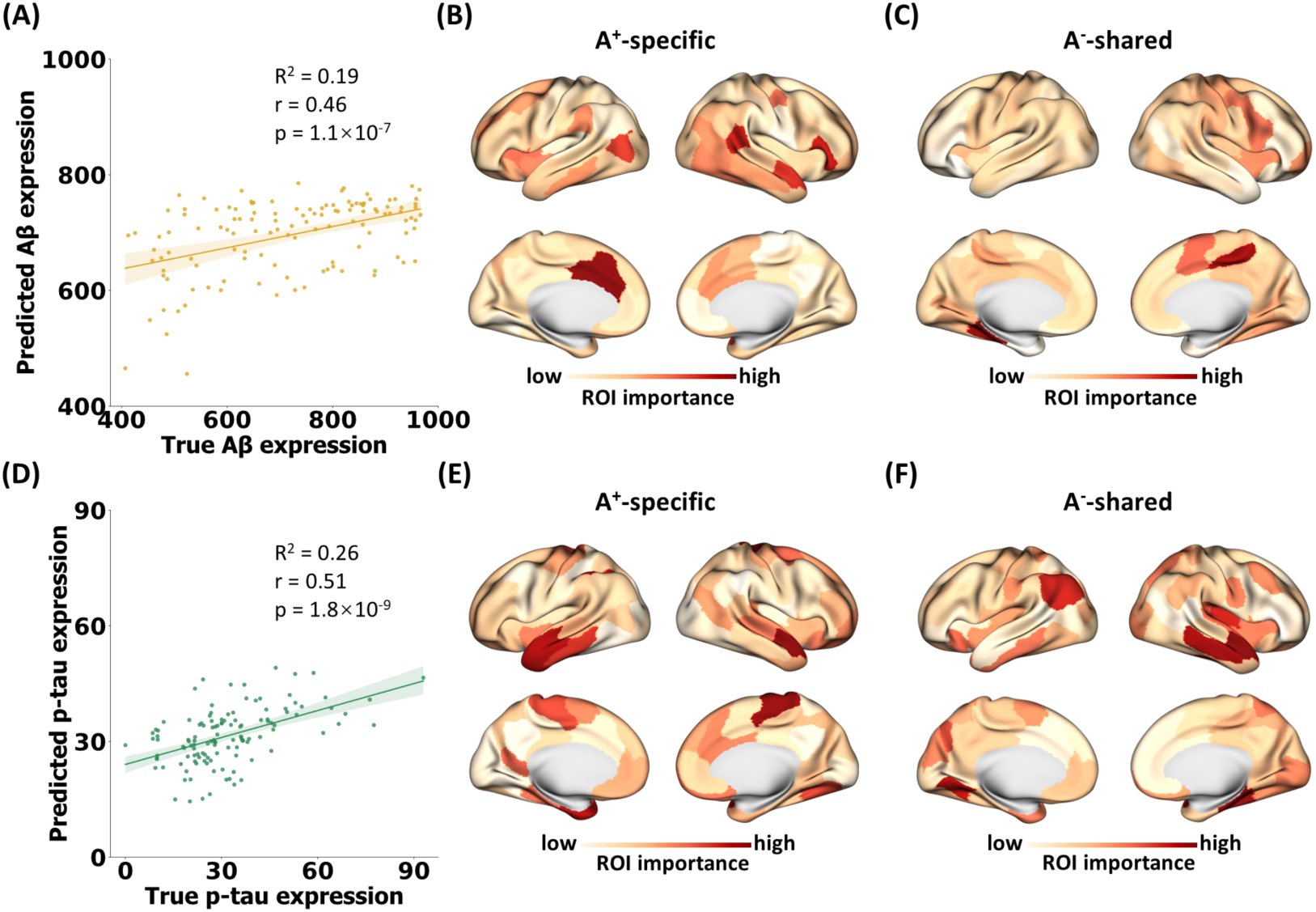
Prediction of individual Aβ and p-tau levels under the A^+^/A^-^ framework using our cGCN model. Prediction accuracy was evaluated across ten repetitions of 5-fold cross-validation. The brain importance was computed by summing the gradients of latent features with respect to input node features of each region. **(A)** The scatter plot between true Aβ expression and predicted Aβ expression averaged from ten 5-fold cross-validated models (R^2^ = 0.19, Pearson’s r = 0.46, p = 1.1 × 10^−7^ based on the one-sided test against the alternative hypothesis that r > 0). **(B)** Brain signature of A^+^-specific dimension for Aβ prediction. **(C)** Brain signature of A^-^-shared dimension for Aβ prediction. **(D)** The scatter plot between true p-tau expression and predicted p-tau expression averaged from ten 5-fold cross-validated models (R^2^ = 0.26, Pearson’s r = 0.51, p = 1.8 × 10^−9^ based on the one-sided test against the alternative hypothesis that r > 0). **(E)** Brain signature of A^+^-specific dimension for p-tau prediction. **(F)** Brain regional signature of A^-^-shared dimension for p-tau prediction.

Subsequently, we defined brain prediction signatures by summing the gradients of latent features with respect to the input brain graphs in both A^+^-specific and A^-^-shared dimensions (See Methods section for details). Larger gradients indicated that functional connections involving the region were more critical for prediction. For Aβ prediction, the A^+^-specific dimension emphasized the gradients involving the bilateral anterior cingulate cortex, right inferior frontal, and superior temporal cortices (Figure 1 (B)). In contrast, the A^-^-shared dimension highlighted gradients involving the left fusiform cortex, right middle cingulate, inferior frontal cortices, and the supplementary motor area (Figure 1 (C)). For p-tau prediction, the A^+^-specific dimension identified critical ROIs with larger gradients in the bilateral paracentral cortices, temporal poles, and the inferior parietal cortex (Figure 1 (E)), while the A^-^-shared dimension emphasized gradients from the right insula, inferior temporal, and left angular cortices (Figure 1 (F)).

To assess the similarity between the four brain signatures shown in Figure 1, we employed Pearson correlation analysis, followed by permutation testing with 1000 iterations to randomly shuffle brain regional importance, which confirmed the statistical significance of the observed correlations.

Notably, a significant correlation emerged only between the A^+^-specific brain signatures for Aβ and p-tau (r = 0.36, p_perm_ = 0.002), primarily driven by overlapping ROIs of high importance, such as the left anterior insula, right superior temporal gyrus, temporal pole, and anterior cingulate cortex.

### Distinctive characteristic profiles between A^+^-specific and A^-^-shared features

To elucidate the phenotyping implications of the A^+^-specific and A^-^-shared latent features as disentangled by our model, we analyzed the brain graphs from the A^+^ group using the well-trained cGCN models to obtain these latent features (Figure S2(A)). We then measured their associations with various characteristic variables (including demographic variables and cognitive measures such as age, MoCA) through representative similarity analysis (RSA)^37^ (see Method section for details) as characteristic profiles. To substantiate the uniqueness of the characteristic profiles of latent features disentangled by our cGCN model, we conducted the same analysis using ablation models devoid of a contrastive module as a control study. We applied the Kruskal-Wallis ANOVA test to examine significant differences in the characteristic profiles related to A^+^-specific, A^-^-shared, and entangled latent features (without contrastive learning). Upon confirming notable differences across those three groups, we conducted Dunn’s post-hoc test to further dissect and compare these distinctions in pairwise groups.

For Aβ prediction, A^+^-specific brain latent features exhibited significant correlations with *APOE* genotypes, total learning ability and learning rate (as assessed by the *Rey Auditory Verbal Learning Test* (RAVLT)) as well as planning and organization skills (as assessed by *Everyday cognition Scale* (ECog)) (Figure 2 (A)). These correlations were markedly stronger than those observed with the A^-^-shared and entangled features. On the other hand, the A^-^-shared latent features showed significant correlations with various cognitive impairment measures such as *Alzheimer’s Disease Assessment Scale-Cognitive 11* (ADAS 11), *Clinical Dementia Rating Scale* (CDR), total scores of *Montreal Cognitive Assessment* (MoCA) and language ability, and forgetting rate of *Rey Auditory Verbal Learning Test* (RAVLT). To examine which cognitive impairment stage contributed most to the identified characteristic profiles, we divided A^+^ subjects into cognitively unimpaired (CU, n = 49), mild cognitive impairment (MCI, n = 47), and dementia (n = 33) subgroups, and performed RSA analyses for each subgroup separately. We observed that in individuals with MCI, the associations between A^+^-specific latent features with learning ability and *APOE*, and between A^-^-shared latent features with ADAS 11, CDR, and MoCA were further increased (Figure S3), compared to the association observed in all A^+^ subjects. This observation suggested that the significant correlations between A^+^-specific, A^-^-shared latent features and these global measures of cognition and function were predominantly derived from MCI subjects. Furthermore, we implemented the RSA within A^+^T^+^ subgroup and figured out that the A^-^-shared latent features were correlated with age and A^+^-specific latent features were correlated with learning ability (Figure S4 (A)).

**Figure 2.**
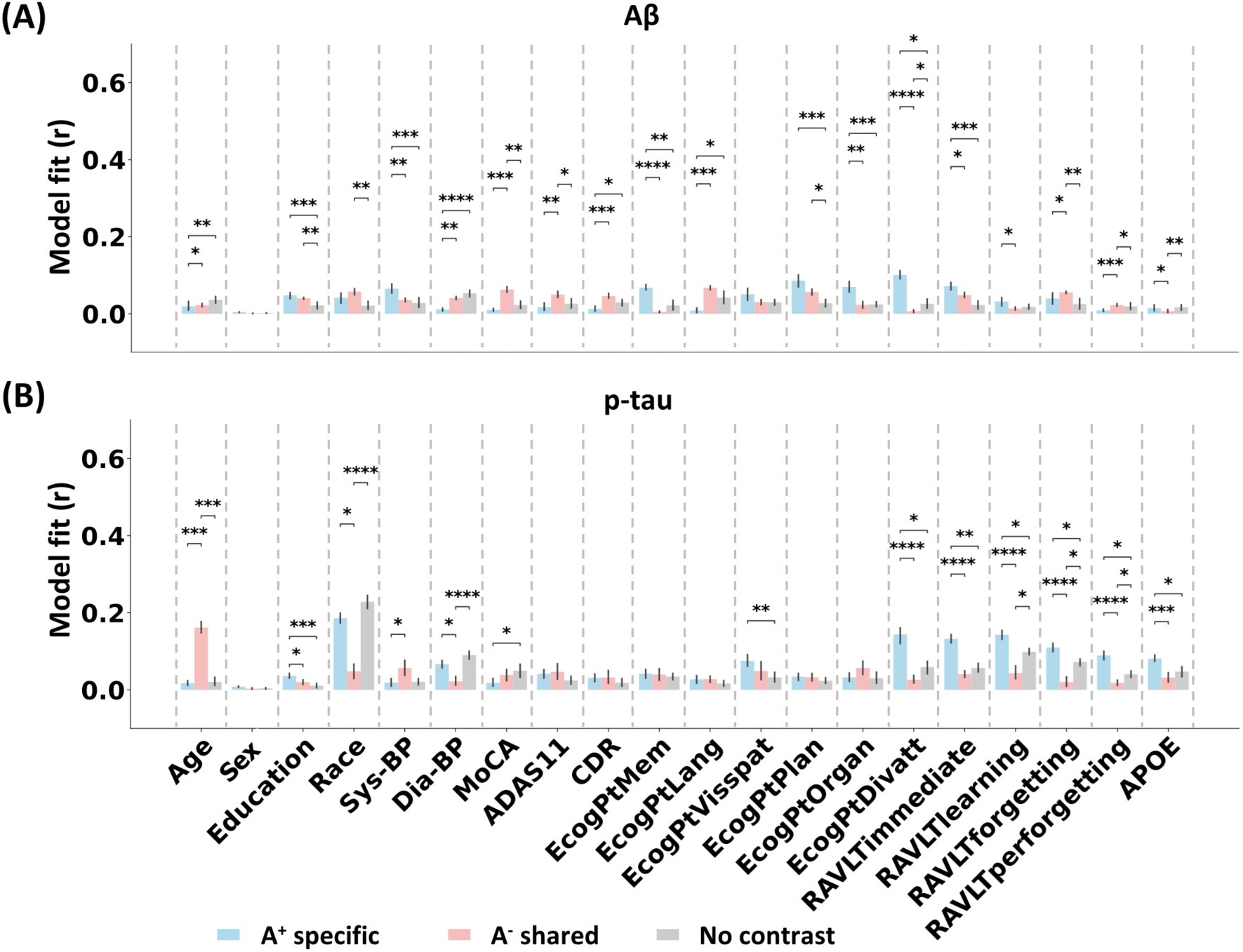
Characteristic profiles of Aβ and p-tau predictive brain latent features in A^+^ subjects. Representative similarity analysis was used to estimate the associations between characteristic variables and latent features, derived from cross-validated models for predicting Aβ and p-tau separately. Data expressed as mean ± s.d. Fisher’s combined probability test confirmed the significance of the correlation between each characteristic variable and latent features from A^+^-specific, A^-^-shared dimensions, and ablation models without a contrastive module in 10 trials. Kruskal-Wallis one-way ANOVA detected group differences in correlations involving latent features disentangled from A^+^-specific, A^-^-shared dimensions, and ablation models without a contrasting module. Dunn’s test was then applied to identify pairwise differences in correlation. False discovery rate (FDR) correction of statistic comparison across all multiple characteristic variables and latent features was implemented. If latent features extracted from the A^+^-specific, A^-^-shared dimensions, and ablation models were significantly correlated with the characteristic variables, and there was a significant difference in pairwise correlations confirmed by Dunn’s test, the significant differences were denoted by *(p ≤ 0.05), **(p ≤ 0.01), ***(p ≤ 0.001), and ****(p ≤ 0.0001). **(A)** Characteristic profiles associated with A^+^-specific, A^-^-shared and entangled brain graphs latent features in predicting Aβ expressions. **(B)** Characteristic profiles associated with A^+^-specific, A^-^-shared and entangled brain graph latent features of subjects in predicting the p-tau expressions. Sys-BP: systolic blood pressure; Dis-BP: diastolic blood pressure; MoCA: Montreal Cognitive Assessment; ADAS11: Alzheimer’s Disease Assessment Scale-Cognitive 11 items; EcogPtMem, EcogPtLang, EcogPtVisspat, EcogPtPlan, EcogPtOrgan, EcogPtDivatt: memory, language, visuospatial ability, planning, organization, dividing attention scores of Everyday Cognition Scale; RAVLTimmediate, RAVLTlearning, RAVLTforgetting, RAVLTperforgetting: evaluation of total learning ability, learning rate, long-term retention and forgetting rate of Rey Auditory Verbal Learning Test.

For p-tau prediction, the A^+^-specific brain latent features showed notable correlations with *APOE* genotypes and several aspects of verbal memory (learning rate, long-term retention and forgetting rate), and attention (Figure 2 (B)). In contrast, A^-^-shared features exhibited a stronger correlation with age and systolic blood pressure. Integrating these results with the characteristic profiles of Aβ predictive latent features revealed that A^+^-specific latent features captured more variations related to learning ability when compared to A^-^-shared latent features. When performing RSA analyses for A^+^ subjects from CU, MCI, and dementia subgroups separately, the correlation between A^+^-specific latent features and *APOE* and verbal memory was notably strong in MCI subgroup. Meanwhile, the correlation between A^-^-shared latent features and age was observed in CU subgroup (Figure S3), suggesting that the association between A^-^- shared latent features and aging was predominantly derived from CU subjects. Additionally, the RSA within A^+^T^+^ subgroup indicated that the A^-^-shared latent features were correlated with age and A^+^- specific latent features were correlated with ability of attention division (Figure S4 (B)).

### Cortical gene expression related to A^+^-specific and A^-^-shared features

To probe the potential molecular pathways underlying the A^+^-specific and A^-^-shared features, we examined the relationship between transcriptomic variations and the identified brain signatures shown in Figure 2. Using the Schaefer parcellation^38^, we extracted transcriptional expression levels of 100 brain regions from the AHBA dataset^14^ and applied PLS regression with a transcriptional expression matrix (100 regions × 15633 genes) to predict the brain signatures to obtain associated genes (Figure S2 (B)) (see Methods section for details). We then conducted gene set enrichment analysis (GSEA) using Metascape (https://metascape.org/gp/index.html#/main/step1)^39^ to analyze the significantly correlated gene sets, aligning with the gene ontology (GO) biological processes, Kyoto Encyclopedia of Genes and Genomes (KEGG), Reactome, and Wiki pathways.

The top 10 genes with significant PLS weights (z-score transformed) associated with the A^+^- specific brain signature predictive of Aβ are shown in Figure 3 (A). Notably, the UBA52 and INTS4P1 genes exhibited high expression levels in the right insula, emerging as the foremost predictive genes.

**Figure 3.**
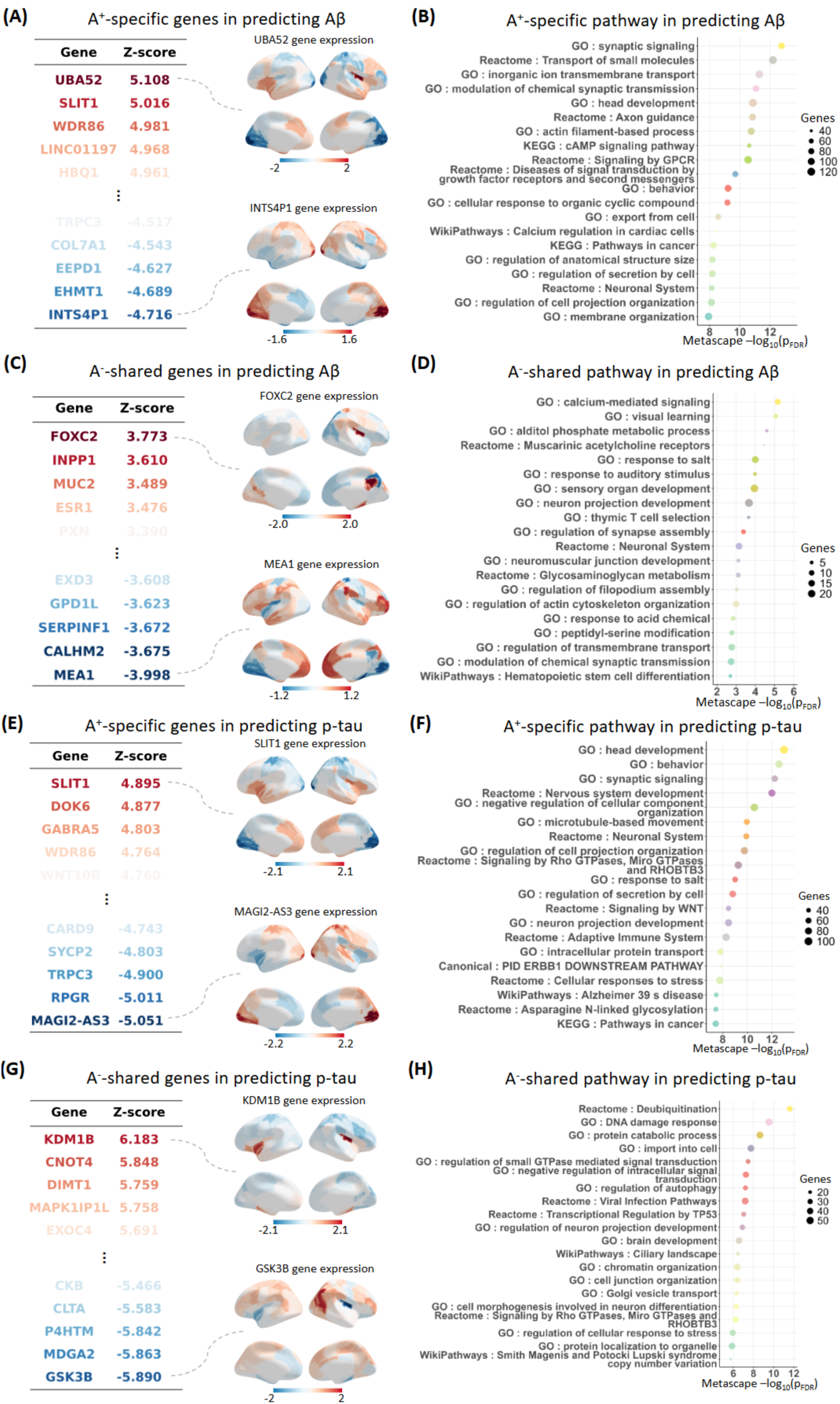
Transcriptomic correlations of Aβ and p-tau predictive brain signatures. PLS analysis was conducted to predict brain signatures (as shown in Figure 1) based on transcriptional expression. Bootstrap analysis with 2000 trials was employed to identify significantly correlated gene sets, with false discovery rate correction applied across all genes. The weight of each gene was divided by its bootstrap standard error to compute the Z score. Gene set enrichment analysis, aligning with gene ontology (GO) biological processes, Kyoto Encyclopedia of Genes and Genomes (KEGG), Reactome, and Wiki pathways, were conducted for the significantly correlated gene sets using Metascape website. **(A), (C), (E), (G)** The top 10 genes with strong positive and negative weights of the first PLS component, showcasing the gene expression level across whole brain regions with the largest PLS positive and negative weights. **(B), (D), (F), (H)** The top 20 enriched functional terms, featuring the bubble plot of the significantly correlated genes for each brain signature. **(A)** Visualization of the most significantly correlated genes of A^+^-specific brain signature in predicting Aβ. **(B)** Bubble plot of the top 20 enriched terms of the significantly correlated genes of A^+^-specific brain signature in predicting Aβ. **(C)** Visualization of the most significantly correlated genes of A^-^-shared brain signature in predicting Aβ. **(D)** Bubble plot of the top 20 enriched terms of the significantly correlated genes of A^-^-shared brain signature in predicting Aβ. **(E)** Visualization of the most significantly correlated genes of A^+^-specific brain signature in predicting p-tau. **(F)** Bubble plot of the top 20 enriched terms of the significantly correlated genes of A^+^-specific brain signature in predicting p-tau. **(G)** Visualization of the most significantly correlated genes of A^-^-shared brain signature in predicting p-tau. **(H)** Bubble plot of the top 20 enriched terms of the significantly correlated genes of A^-^-shared brain signature in predicting p-tau.

UBA52, which encodes ubiquitin, is crucial for eliminating insoluble aggregates like Aβ plaques and neurofibrillary tangles through the ubiquitin-dependent protein degradation system^40^. Impairment of this system may prevent the reduction of insoluble pathogenic fragments in the brain. Further investigations into the molecular functions of the correlated genes revealed pathways central to synaptic signaling, transport of small molecules, modulation of chemical synaptic transmission, head and axon development, and the cAMP signaling pathway (Figure 3 (B), S5 (A)). In relation to the A^-^-shared brain signature, the FOXC2 gene was overexpressed in the insula and posterior cingulate cortex, while the CALHM2 gene was overexpressed in the lingual gyrus and bilateral medial orbitofrontal cortices (Figure 3 (C)). The primary biological functions linked to these genes encompassed pathways involved in calcium-mediated signaling, visual learning, auditory stimuli, and neuron projection development (Figure 3 (D), S4 (B)).

For the p-tau prediction, A^+^-specific brain signature was associated with genes such as SLIT1 and MAGI2-AS3, which were predominantly overexpressed in the primary visual and paracentral cortices (Figure 3 (E)). This signature was linked to molecular pathways pertinent to behavior, synaptic signaling, and microtubule-based movement (Figure 3 (F), S4(C)). In contrast, genes such as KDM1B and GSK3B, significantly correlated with the A^-^-shared brain signature, were overexpressed in the insula and middle temporal and occipital cortex (Figure 3 (G)). These genes were linked to molecular pathways such as deubiquitination, DNA damage response, protein catabolic processes, import to cell, and autophagy regulation (Figure 3 (H), S4 (D)). For example, GSK-3β is a central kinase involving the regulation of autophagy^41–43^, with its dysregulation leading to key disease characteristics such as tau hyperphosphorylation, and impaired memory, neurogenesis, and synaptic function^44–46^.

To delve deeper into the potential intersections between our identified gene sets and AD risk genes, we aligned the gene sets identified through PLS analyses with gene loci associated with AD risk from a previous genome-wide association study (GWAS)^47^ meta-analysis. We evaluated the concordance between gene sets correlated with the different dimensions of Aβ and p-tau and those implicated in GWAS using the rank biased overlap^48,49^ (a robust measure for assessing non-conjoint lists). This comparison revealed a pronounced concordance between gene sets correlated with Aβ-predictive and p-tau-predictive A^+^-specific dimensions and previous GWAS discoveries (Figure S6 (A, B)). In contrast, the gene sets correlated with A^-^-shared dimensions showed limited similarity to the AD risk GWAS results. In addition, a multi-gene-list meta-analysis was conducted to ascertain overlapping molecular pathways reflected by these gene sets^39^. Our results highlighted a set of common pathways across all five gene sets, including synaptic signaling, neural systems, neuron projection development, and the modulation of chemical synaptic transmission (Figure S6 (C, D)). Notably, compared to the A^-^-shared dimension, these pathways were more prominently expressed in the gene sets correlated with the A^+^- specific dimension, aligning closely with findings from GWAS of AD^47^. Specifically, the associations between A^+^-specific dimensions linking to Aβ and p-tau and pathways involving synaptic signaling were enhanced.

### Transcriptional signatures linked to canonical cells in the central nervous system

Previous studies have underscored the critical role of cellular diversity, inter-cell interactions, and transcriptional abnormalities in the development of AD^33,34,50^. To further explore the associations between transcriptional expression patterns of central nervous system cell types and Aβ and p-tau expressions, we categorized the gene lists identified from PLS analyses into seven canonical cell categories: excitatory neurons, inhibitory neurons, microglia, endothelial cells, oligodendrocytes, astrocytes, and oligodendrocyte precursors (OPCs)^51^. We then quantified the gene expressions involving different molecular activities of these cell types and associated them with brain signatures of different dimensions (Figure 4). We observed that genes and molecular pathways linked to A^+^-specific signatures were associated with a wider variety of cell types compared to A^-^-shared signatures. For instance, the Aβ- linked A^+^-specific dimension was correlated with molecular pathways including cell activation, TYROBP causal network, actin cytoskeleton organization and cAMP signaling related to the microglia, oligodendrocytes, and inhibitory neurons. Similarly, in p-tau prediction, compared to A^-^-shared dimension, the A^+^-specific dimension was involved in enriched molecular pathways related to synaptic signaling, membrane potential, cellular anatomical entity morphogenesis, and plasma membrane-bounded cell projection organization, associated with microglia, astrocytes, and inhibitory neurons. Furthermore, when comparing cell type-specific molecular pathways associated with the A^+^-specific dimension predictive for Aβ and p-tau, both sets of pathways were involved in synaptic signaling and various processes related to neurotransmitter transport and neuronal development.

**Figure 4.**
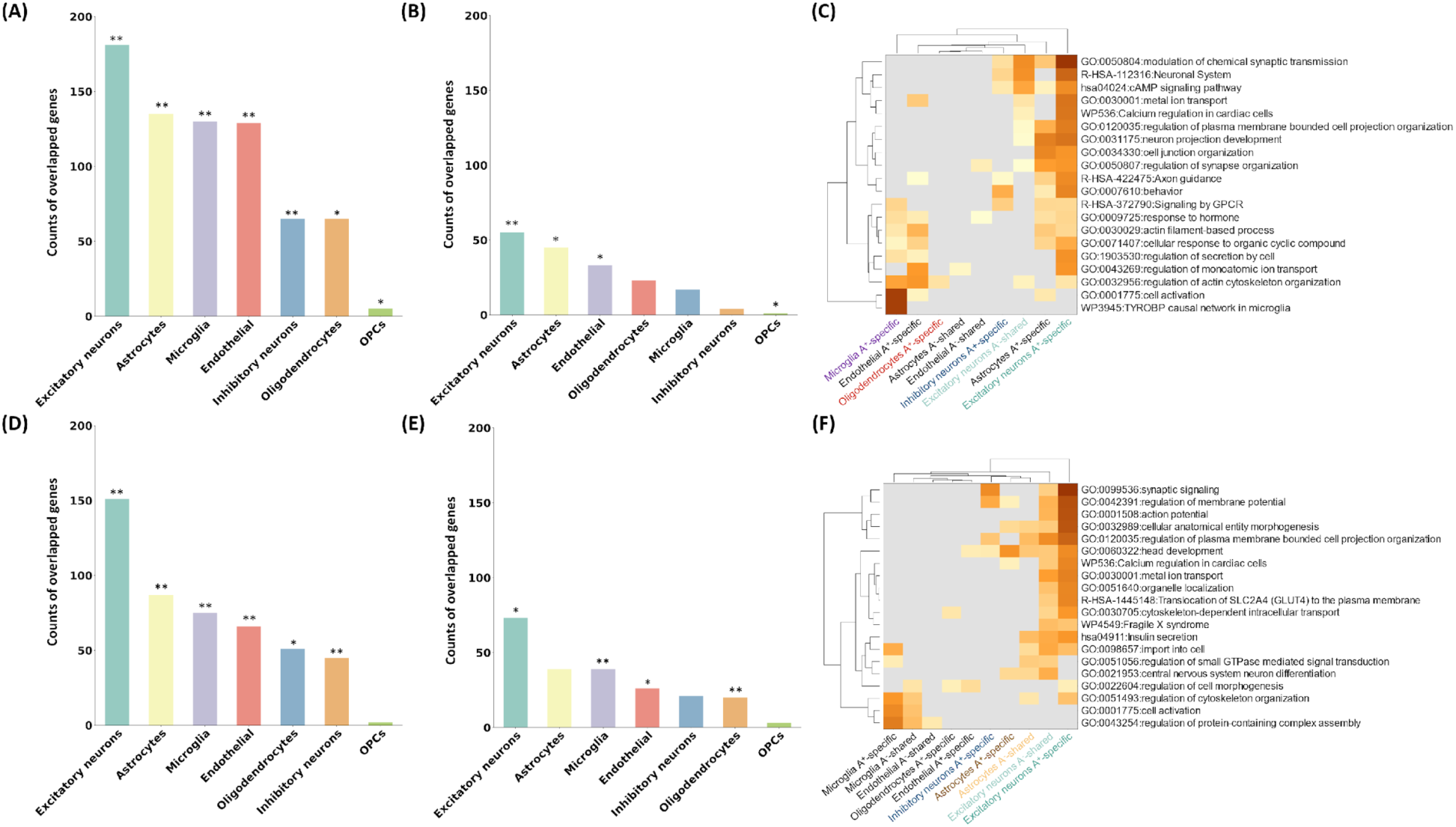
Cell type-specific transcriptomic expression related to Aβ and p-tau proteins. The significance of overlapping counts between Aβ and p-tau predictive gene lists and cell-specific gene lists was confirmed by 1000 permutation test. It was annotated by * (p ≤ 0.05), ** (p ≤ 0.01). **(A)** Counts of genes of A^+^-specific dimension for Aβ prediction, related to each cell type. **(B)** Counts of genes of A^-^-shared dimension for Aβ prediction, related to each cell type. **(C)** Top 20 enriched molecular pathways of significantly overlapping genes of each cell type linked to Aβ expression. **(D)** Counts of genes of A^+^-specific dimension for p-tau prediction, related to each cell type. **(E)** Counts of genes of A^-^-shared dimension for p-tau prediction, related to each cell type. **(F)** Top 20 enriched molecular pathways of significantly overlapping genes of each cell type linked to p-tau expression.

## Discussion

To our knowledge, this study represents a pioneering effort to elucidate the heterogeneity of Aβ and p-tau using whole-brain functional graphs. Although some interpretations of our findings may be speculative, they serve as a foundation for enhancing our understanding of the complex nature of AD pathological proteins. We designed a novel cGCN deep learning model to disentangle variations of CSF Aβ and p-tau into A^+^-specific and A^-^-shared dimensions. Our findings revealed that A^+^-specific brain latent features were more predictive of individual Aβ and p-tau levels compared to the features intertwined with A^-^. Furthermore, our analyses uncovered distinct associations between A^+^-specific brain latent features and characteristic variables (age, RAVLT, etc.), gene expression patterns, and molecular pathways, as well as associations between A^-^-shared latent features and these measures. Importantly, A^+^- specific latent features predictive of both Aβ and p-tau expression show correlations with cognitive functions such as attention and memory, as well as molecular pathways encompassing synaptic signaling, neuron systems, and behavior. Additionally, the identified A^-^-shared latent features in predicting p-tau correlated to the aging-related pathway.

### The neuropsychological profiles associated with Aβ and p-tau prediction

The A^+^-specific and A^-^-shared dimensions linked to Aβ highlighted the influence of Aβ pathology on different cognitive functions. The A^+^-specific dimension, marked by brain network connections involving the bilateral anterior cingulate cortex, right inferior frontal and superior temporal cortices, correlated strongly with episodic memory, planning and organization. Aligning with previous findings that these areas have been affected by higher neurofibrillary tangles^52–55^, A^+^-specific dimension emphasized the executive dysfunction observed in Alzheimer’s continuum. On the other hand, the A^-^- shared dimension, dominated by connections in the left fusiform gyrus and right middle cingulate, correlates with global measures of cognition and function such as CDR. Additionally, when examining the MCI group, the correlation between this dimension and the CDR was enhanced. Previous studies have indicated that functional connectivity alternations in these dominant regions of the A^-^-shared dimension correlated with CDR severity in MCI^56,57^. In line with this observation, the correlation between this dimension and CDR derived from MCI groups highlights its potential application for predicting cognitive impairment severity within MCI.

In addition to Aβ pathology, the A^+^-specific and A^-^-shared dimensions offer novel insights into p-tau pathology. The A^+^-specific dimensions that were derived from brain network features in the bilateral paracentral cortices, temporal pole, and inferior parietal cortex, predicted p-tau levels and strongly correlated with learning capabilities such as learning rate and long-term retention. In AD patients, abnormal connections and metabolic activity in these brain regions are associated with dysfunctional word learning^58^, memory consolidation^59,60^, and semantic processing^61,62^. Therefore, the A^+^-specific dimension may depict aspects of p-tau pathology involving learning-related cognitive functions^63,64^. On the other hand, the A^-^-shared dimensions, involving brain network features in the right insula and left angular cortex, showed a strong correlation with age. Previous fMRI studies suggest that compared to younger normal adults, in elderly people, the right insula is activated during affect induction and perception tasks^65^, while gray matter in the left angular gyrus is reduced^66^. Therefore, the A^-^-shared dimension may capture aspects of normal aging, supporting the view that mild tauopathy can be found in healthy aging^67^.

### Molecular pathways unveiling new insights into pathology of Aβ and p-tau

The complexity of Aβ pathology is manifested through an intricate interplay of various molecular pathways. First, the GSEA of their brain signatures both reveal the importance of molecular pathways related to the neuronal system, synaptic activities, and Ca^2+^ regulation. This finding aligns with the foundational amyloid cascade and oligomerization hypotheses, emphasizing the central roles of synaptic dysfunction and Ca^2+^ signaling disturbances in Aβ pathology^68–71^. However, A^+^-specific and A^-^-shared dimensions diverge in other pathways. The A^+^-specific dimension highlighted the role of neural calcium homeostasis in Aβ pathology, focusing on pathways involved in inorganic ion transmembrane transport, cAMP signaling, and G protein-coupled receptor (GPCR) signaling, which are both closely linked to Ca^2+^ regulation^72^. In contrast, the A^-^-shared dimension emphasized downstream consequences of Aβ pathology or co-pathology, such as the pathways related to visual learning, muscarinic acetylcholine receptor function, and auditory responses. The impaired function of the muscarinic acetylcholine receptor, linked to abnormal processing of amyloid precursor protein^73^, plays a pivotal role in both auditory and visual processing realms^74,75^, which are crucial for language processing^76,77^. Combining the correlation between A^-^-shared dimension and language function, this dimension might emphasize the impact of Aβ pathology on language dysfunction through pathways involving the muscarinic acetylcholine receptor. Moreover, the dysfunction of muscarinic acetylcholine receptor and visual learning pathways is typically associated with auditory and visual hallucination observed in patients with Lewy body AD^78–81^. Thus, the A^-^-shared dimension might also capture some variations of Lewy body copathology in AD.

The GSEA outcomes for p-tau revealed unique pathways. The A^+^-specific dimension highlighted pathways linked with neuroinflammation, which is typically observed in later stages of AD progression^82^.

Specifically, pathways involving GTPase and WNT signaling were related to neuroinflammation processes^83,84^. Additionally, this dimension highlighted the molecular pathways related to astrocyte activity. Astrocytosis and neuroinflammation are both associated with elevated expression levels of glial fibrillary acidic protein (GFAP), which contributes to propagation of tau^85,86^. Our A^+^-specific dimension might serve as indirect evidence to support the association between GFAP and tau pathology^87^.

Conversely, the A^-^-shared dimension highlights pathways implicated in the normal aging process, including deubiquitination^88^, DNA damage response^89^, protein catabolism, autophagy regulation^90^ and GSK-3β function. Consistent with our earlier discussions on the neuropsychological profile of the A^-^- shared dimension, these molecular pathways underscored the overlap between this dimension and normal aging in biological processes.

### Overlapping profiles between Aβ and p-tau linked A^+^-specific dimensions

As expected, the brain signatures identified from the A^+^-specific dimensions for predicting Aβ and p-tau showed a significant correlation, originating from overlapping regions, such as the left fusiform gyrus, left posterior insular, right superior temporal cortex and temporal pole. These regions are implicated in the progressive distribution of tau aggregations in the Braak stages III-IV^91,92^, along with marked increases in cognitive impairment^93,94^. From these stages onward, the correlation between Aβ and tau deposition becomes particularly significant^95^. Integrating these findings, the correlation of A^+^-specific brain signatures in predicting Aβ and p-tau might reflect the underlying neurobiological characteristics of these Braak stages. Furthermore, our GSEA revealed that A^+^-specific dimensions linked to Aβ and p-tau both involved molecular pathways related to synaptic regulation, with a pronounced involvement of excitatory neurons. This was congruent with the observation that synaptic dysfunction, especially the dysregulation of excitatory synapse, serves as a hallmark of Aβ and p-tau pathology^96,97^. Additionally, the enrichment of genes associated with the functions of astrocytes and microglia in the A^+^-specific dimensions aligns with current knowledge that the abnormal tau oligomers coincide with excessive synaptic engulfment by these glial cells^98^.

## Limitations and future research directions

Several limitations in this study warrant consideration. First, the absence of an independent cohort to validate the biological findings from our model raises the need for future research with larger cohorts, incorporating fMRI and CSF biomarkers to assess the reproducibility of our results. Second, the growing focus on alternative modalities for in vivo detection of Aβ and p-tau, such as plasma and positron emission tomography^99^, prompts the exploration of diverse and shared biological processes captured between these modalities and CSF in this study. Third, this study focused on the biomarker interpretation of A^+^/A^-^ contrastive framework. However, the shared neuropathology between A^+^ and A^-^, whether originating from A^-^T^-^CU or A^-^T^-^MCI, remains underexplored. In the future, it would be beneficial to include more subjects categorized as A^-^T^-^CU in the background group to further clarify this question. Fourth, the gene expression data from the AHBA sampled from six healthy donors without a diagnosis of Alzheimer’s continuum is susceptible to inter-subject variability and changes specific to aging and Alzheimer’s continuum. This limits the examination of transcriptome–neuroimaging associations across groups. Despite these limitations, the statistical methods used in this study linking brain signatures to gene expressions have shown to be statistically robust^17,49^ and have biological plausibility. Future endeavors should involve gene expression arrays from independent postmortem brains from demographic-matched individuals to reaffirm the gene enrichment findings associated with CSF Aβ and p-tau.

In summary, our study advanced a novel cGCN-based deep learning framework that adeptly isolates brain functional variations associated with CSF Aβ and p-tau into A^+^-specific and A^-^-shared dimensions. The functional connectivity features from right superior temporal and anterior cingulate cortices of A^+^-specific dimension contributed mostly to predicting individual Aβ and p-tau levels, reflecting the Aβ and p-tau pathology in A^+^ subjects. The Aβ-predictive A^-^-shared dimension was associated with language processing and the p-tau-predictive A^-^-shared dimension was associated with normal aging processes. Linking these brain signatures with gene expression data, we observed that the molecular pathways involving synaptic dysfunction played a key role in Aβ and p-tau pathologies.

Moreover, the pathways involved in metabolic activities of excitatory neurons, astrocytes, and microglia, represent core molecular-biological functions associated with Alzheimer’s continuum. Overall, our findings enrich the comprehension of AD’s neurodegenerative processes across neurobiological, neuropsychological and molecular aspects and offer a model for potential application to disentangle the variations of disorders with shared comorbidity.

## Methods

### ADNI dataset

The Alzheimer’s Disease Neuroimaging Initiative (ADNI) is a longitudinal observational program that enrolled participants aged 55 to 90 years to collect and analyze biomarkers related to the progression of AD^27^. Launched in 2003, ADNI has collected serial MRI and biological markers such as CSF Aβ and p-tau proteins. Participant recruitment for ADNI was approved by the Institutional Review Board of each participating site. In this study, ADNI phase 1, phase GO/2 and phase 3 were used.

Exclusion criteria included current use of psychoactive medication, a history of schizophrenia, substance use and various mental disorders (More details can be found in literature^27^). A total of 161 participants (83 female [52%]; mean [std] age, 72 [79-65] years) who had both fMRI and CSF data available were utilized in this study (Table S2).

### PREVENT-AD dataset

The Pre-symptomatic Evaluation of Experimental or Novel Treatments for Alzheimer’s Disease (PREVENT-AD) program recruited cognitively healthy participants over 55 years old with a first-degree family history of AD in Canada. The primary objective of this program is to facilitate the discovery of safe and promising pharmacological interventions that can slow down the progression of pre-symptomatic AD by analyzing biomarkers from diverse modalities. Exclusion criteria for participants included cognitive disorders and the use of acetylcholinesterase inhibitors. More details about this dataset can be found in literature^28^. A total of 128 participants (86 female [67%]; mean [std] age, 63 [58-68] years) who had both fMRI and CSF data available were utilized in this study (Table S3).

### Characteristic information

Demographics including age, sex, education year, race, systolic blood pressure, and diastolic blood pressure were analyzed. To assess dementia symptoms, various measurements were employed: *Montreal Cognitive Assessment* (MoCA)^100^, *Alzheimer’s Disease Assessment Scale-Cognitive 11*(ADAS 11)^101^, *Clinical Dementia Rating Scale* (CDR)^102^. Additionally, the *Everyday Cognition Scale* (ECog)^104^ evaluated multidimensional functions in older adults, focusing on everyday functions across six cognitive domains: memory, language, visuospatial abilities, planning, organization, and divided attention. The *Rey Auditory Verbal Learning Test* (RAVLT) served as a crucial tool for assessing verbal memory of participants^105^. The RAVLT involved presenting participants with a list of 15 unrelated words to recall over five learning trials. Following an interference word list and a delay, participants were tasked with recalling the initial word list. Various summarized scores derived from raw RAVLT scores provided nuanced insights into distinct cognitive processes^106,107^. These include RAVLT Immediate (the sum of scores from 5 first trials) highlighting total learning ability; RAVLT Learning (the score of trial 5 minus trial 1) emphasizing learning rate; RAVLT Forgetting (the score of trial 5 minus the score of the delayed recall) highlighting long-term retention; RAVLT Percent Forgetting (RAVLT Forgetting divided by the score of trial 5) emphasizing forgetting rate or delayed memory. To be noticed, only CDR assessment was available between ADNI and PREVENT-AD dataset, other assessments were available for ADNI dataset.

### CSF protein collection and processing

In the ADNI dataset, CSF was collected via lumbar puncture using either a 2- or 24-gauge spinal needle, with subsequent storage in tubes. The quantification of Aβ_1–42_ and p-tau_181_ in CSF aliquots was conducted using the multiplex xMAP Luminex platform (Luminex Corp, Austin, TX) with Innogenetics (INNO-BIA AlzBio3; Ghent, Belgium; for research use–only reagents) immunoassay kit–based reagents, as described in previous studies^108^. The Innogenetics kit included monoclonal antibodies specific to Aβ_1–42_ (4D7A3) and p-tau_181_ (AT270), chemically attached to uniquely coded beads, along with analyte-specific detector antibodies (HT7, 3D6). *APOE* genotyping was measured using EDTA blood samples and TaqMan quantitative polymerase chain reaction assays. Genotyping covered *APOE* nucleotides 334 T/C and 472 CT, using an ABI 7900 real-time thermocycler (Applied Biosystems, Foster City, CA), with freshly prepared DNA from EDTA whole blood.

In the PREVENT-AD dataset, lumbar punctures utilized a Sprotte 24-gauge atraumatic needle^109^. CSF samples underwent centrifugation within four hours to exclude cells and insoluble material. The CSF AD biomarkers p-tau_181_ and Aβ_1-42_ were measured using validated Innotest ELISA kits from Fujirebio, following procedures from the BIOMARKAPD consortium^110^. *APOE* DNA extraction from 200 μl whole blood was conducted using a QIASymphony apparatus and the DNA Blood Mini QIA Kit (Qiagen, Valencia, CA, USA), adhering to the manufacturer’s instructions^28^.

### Resting-state fMRI acquisition and preprocessing

For the ADNI dataset, resting-state fMRI was acquired from 3T Philips scanners. For the PREVENT-AD dataset, resting-state fMRI was scanned using gradient echo planar imaging pulse sequences on Siemens TIM Trio 3T MRI scanners. Detailed MRI scanning procedures for both datasets can be found in literature^27,28^.

All the fMRI data were preprocessed using the established fMRIPrep pipeline^111^. The T1 weighted image was corrected for intensity nonuniformity and then skull was stripped as T1w reference. Spatial normalization was done through nonlinear registration with the T1w reference^112^. The brain tissue (cerebrospinal fluid, white matter, and grey matter) was segmented from the T1w reference using FAST (FSL)^113^. The BOLD reference was then transformed to the T1w reference using a boundary-based registration method, configured with nine degrees of freedom to account for distortion remaining in the BOLD reference^114^. Head-motion parameters (six rotation and translation parameters of volume-to-reference transform matrices) were estimated with MCFLIRT (FSL)^115^. BOLD signals were slice-time corrected and resampled onto the participant’s original space with head-motion parameters, susceptibility distortion correction, and then resampled into standard space, generating a preprocessed BOLD run in MNI152NLin2009cAsym space. Automatic removal of motion artifacts using independent component analysis (ICA-AROMA)^116^ was performed on the preprocessed BOLD time-signals in MNI space after removal of non-steady-state volumes and spatial smoothing with an isotropic Gaussian kernel of 6 mm FWHM (full-width half-maximum).

### Calculation of functional connectivity

The preprocessed fMRI was averaged into time series of 100 regions of interest (ROIs) defined by the Schaefer parcellation^38^. Pearson correlation was computed between the time series of each pair of ROIs, resulting in a 100 × 100 FCs matrix for each participant. Fisher’s r-to-z transformation was applied to enhance the normality of connectivity, followed by z-score normalization.

### Contrastive graph convolutional network model

#### Graph convolutional network (GCN)

Utilizing a brain functional network framework, where each brain region serves as a node, the goal of GCN^117^ is to derive features in a latent space for various prediction tasks concerning each subject. Following established functional graph construction steps^118,119^, an undirected brain functional graph is defined as *G* = (***V***, ***E***). *E* ∈ *R*^M×M^ represents the adjacency matrix constructed from thresholded and binarized functional connectivity (Figure S1), where *M* is the number of ROIs. The node feature matrix *V* ∈ *R*^M×M^ is constructed from original functional connectivity coefficients, with each element representing Pearson’s correlation coefficients of pairwise ROIs. The core idea of GCN involves utilizing convolutional operators to iteratively aggregate information from neighboring nodes^120^. This enables GCN to capture both local and global relationships across ROIs within the brain graph, thereby extracting meaningful representations of brain regions.

In our study, we opted for a dynamic graph convolutional operator over the conventional one, given its demonstrated superiority in our prior study on brain dysfunction in attention-deficit/hyperactivity disorder^121^. During training, the dynamic graph convolutional operator dynamically calculated the adjacency matrix based on latent features of each ROI and aggregated the latent features of top-*k* neighboring ROIs. This approach enhances model generalizability by introducing sparse regularization through the aggregation of latent features from *k* brain regions. More details about the dynamic graph convolutional operator can be found in our previous study^121^.

#### Contrastive learning

Contrastive learning is an emerging technique to disentangle target-specific features from those shared with the background^22,122^. This approach has been integrated into a contrastive variational autoencoder network, comprising two variational encoders with similar structures: one encoder trained solely on the target dataset and another trained on a combination of target and background datasets. The objective is to maximize the divergence in the distribution of latent features encoded from the target and background datasets while ensuring these latent features can reconstruct the original features. In our study, the target dataset consisted of A^+^ subjects and the background dataset consisted of A^-^ subjects, with subjects having CSF Aβ_1-42_ levels below 976.6 pg/ml^35,36^ labeled as A^+^ and others as A^-^.

Recent studies have shifted from traditional contrastive learning focused on reconstruction to maximizing mutual information between encoded latent features and the original features for downstream graph prediction tasks^123,124^. This approach has shown promising results in various biological applications, such as cell clustering and epilepsy diagnosis^125,126^. Therefore, we adopted a contrastive learning framework derived from deep graph informax^124^, utilizing a binary cross-entropy loss to discriminate latent features encoded from the A^+^ group and the combination of A^+^ and A^-^ groups. The contrastive loss is defined as: *L_c_* = −log(*f_D_*(***H***^(*f*)^, ***S***^(*f*+1)^)) − log(1 – *f*_D(H_*^(f)^*_, **S**_^(f+1)^_))_ (2), where ***H***^(*f*)^ and ***H***^(*f*)^ denote the locally representative features encoded in the convolutional layer from the A^+^ group and the combination of A^+^ and A^-^ groups, respectively. ***S***^(*f*+1)^ represents the globally representative features summarized from the readout layer (the hidden layer next to the final convolutional layer), which transforms the high-dimensional feature matrix from the A^+^ group into a low-dimensional feature vector. The discriminative function *f*_$_ produces probability scores to assess the similarity of latent features of A^+^ group encoded by the A^+^-specific and A^-^-shared encoders. More details about the theory of this contrastive loss can be found in the formula appendix in references^123,124^.

#### Model training guided by CSF Aβ and p-tau prediction

To verify our assumption that A^+^-specific latent features would more accurately capture the variance of AD pathology in the A^+^ group as opposed to the normal aging-related variance in the A^-^ group, we aimed to use these features to predict two critical neuropathological proteins (Aβ and p-tau), key biomarkers for AD screening^3^. Utilizing baseline CSF protein and fMRI data from 289 subjects across the ADNI and PREVENT-AD cohorts, we first pretrained the model with contrastive loss to extract A^+^-specific and A^-^-shared representative features from the two encoders. In the finetuning step, we employed a root mean square error loss to minimize the distance 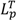 between the true and predicted neuropathological proteins in the A^+^ group, and 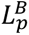 for the A^-^ group. The overall loss during model training in the finetuning step is expressed as: 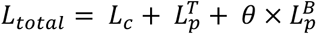, where θ is a parameter controlling the predictive power in the A^-^ group. Predictions for CSF Aβ_1-42_ and p-tau_181_ expression levels (pg/ml) were performed separately.

We validated the model performance through ten repetitions of 5-fold cross-validation on the A^+^ group, given its greater clinical significance compared to the A^-^ group^127^. Prediction results were quantified by computing the R-squared (R^2^) value and Pearson correlation coefficient between the observed protein expressions and predicted ones, averaged across ten repetitions. To interpret the identified predictive signatures, we calculated sum of the gradients for all subjects according to literature^128^, with each gradient representing the differential of predicted values with respect to the input brain graphs.

### Comparing our model with other typical models

To demonstrate the superiority of our model in predicting neuropathological proteins, we compared its performance with several well-established machine learning algorithms, including Ridge, Lasso, support vector regression, XGBoost, random forest, and multi-layer perceptron (MLP) regression models. We calculated the R^2^ between observed and predicted protein expressions using the same cross-validated as described above (Aβ prediction: 0.15 ± 0.02 and p-tau prediction: 0.16 ± 0.02) (Figure S7). We then used Wilcoxon signed-rank tests to statistically compare the R^2^ values of our model against those of the next best-performing method, affirming its significant superiority (Aβ prediction: Z = 2.15, p = 0.03; p-tau prediction: Z = 2.23, p = 0.02) (Figure S7). The higher R^2^ values in scatterplots of our model (Figure 1) were attributed to the ensemble learning effect^129,130^, combining the prediction results of models derived from multiple repetitions of cross-validation.

To systematically examine the influence of different model modules on prediction accuracy, we performed an ablation study by iteratively removing key modules: the graph convolutional operator, the pretraining phase, and the contrastive module (Figure S8). The results indicated that all these modules enhanced the model performance, with the graph convolutional operator being the most critical.

Moreover, to assess the impact of demographic differences between the A^+^ and A^-^ groups on our contrastive learning framework, we performed random subsampling of A^-^ subjects, matching them to A^+^ based on age and sex. After retraining the models, the prediction performance showed no significant difference (Figure S9) (Aβ prediction: Z = -0.77, p = 0.45; p-tau prediction: Z = -0.76, p = 0.46).

Additionally, we investigated the influence of cohort site differences on prediction performance using partial correlation analysis, which controlled for site labels when assessing the relationship between predicted and true scores. As shown in Figure S10, no significant difference was observed (Aβ prediction: Z = -0.08, p = 0.94; p-tau prediction: Z = -0.53, p = 0.60).

### Model performance under different contrastive frameworks

To validate the robustness of our models across different contrastive strategies under the AT/N framework, we re-defined the target group (n = 248) as individuals characterized by either positive Aβ (A^+^T*, Aβ < 976.6 pg/ml) or positive p-tau (A*T^+^, p-tau > 21.8 pg/ml) ^131^ and the background group as A^-^ T^-^ individuals (N=41). Demographic information for these subjects is summarized in Table S4. Due to the severe imbalance in sample sizes between A^-^T^-^ and the combination of A^+^T* and A*T^+^, we applied 10 times augmentation to the functional connectivity for the A^-^T^-^ group, details of which can be found in reference^132^. In short, we used slide windows to randomly select subsequences of time signals to construct augmented FC in A^-^T^-^ group. We trained the models for predicting the CSF Aβ and p-tau expression while contrasting the latent features of target group with all subjects. The results show that our models outperformed other approaches in prediction of Aβ and p-tau expression of individuals within the A^+^T* and A*T^+^ groups (Figure S11).

To confirm the advantage of contrastive strategies under the AT/N framework, we conducted a control analysis using the traditional diagnostic framework. We contrasted the target group, consisting of dementia and MCI patients, against all subjects including CU ones. Neither conventional machine learning models nor our proposed model produced generalizable predictions in the target group (Figure S12).

### Contrastive effect exploration

To confirm the efficacy of our model in disentangling latent features between the A^+^ and A^-^ groups, we employed the Jensen–Shannon divergence to measure the distances of latent features within the A^+^ group and those between the A^+^ and A^-^ groups. We then applied two-sample t-tests to assess the statistical significance of these distances. Under the contrastive learning framework, the latent features showed significantly smaller inter-subject distances within the A^+^ group compared to those between the A^+^ and A^-^ groups (Figure S13 (A)), with Cohen’s d of -8.09 for Aβ prediction and -7.49 for p-tau prediction, both with p values < 0.0001. This difference became non-significant when contrastive module was removed from our model. To further investigate the model’s efficacy in disentangling graph attributes, we calculated three critical graph metrics (clustering coefficient, modularity, and path length) using latent features obtained from our models and compared them between A^+^ and A^-^ subjects. We also calculated these metrics using raw functional connectivity as a baseline. As shown in Figure S13 (B), our cGCN model distinguished graphs in latent space between A^+^ and A^-^ subjects, outperforming both the baseline set by raw functional connectivity and the ablation GCN model without the contrastive module. Additionally, to demonstrate the A^+^-specific encoder capture the A^+^-specific variations predictive for Aβ and p-tau expression of A^+^ subject, we used the well-trained A^+^-specific encoder to predict Aβ and p-tau expression of A^-^ subject. As shown in Figure S14, the A^+^-specific encoder was not able to predict neuropathological proteins expression in A^-^ subjects.

To investigate the reasons for the inferior predictive performance of our model under the traditional diagnostic framework, we calculated the Pearson correlation between each connectivity feature and neuropathological proteins for subjects in the target and background groups under different contrasting frameworks. We then counted the functional connections uniquely correlated to neuropathological proteins in the target groups or inversely correlated to neuropathological proteins between the target and background groups (Figure S15). Our results indicated a higher incidence of functional connections specifically correlated to neuropathological proteins in target groups defined using AD neuropathological biomarkers compared to groups defined by clinical symptomatology. These findings suggest that the AT/N biomarker framework is more effective in elucidating variations in neuropathological proteins than traditional clinical diagnoses, underscoring the efficacy of a biomarker-oriented approach in advancing our understanding of Alzheimer’s continuum neuropathology.

### Characteristic profile obtained using representative similarity analysis

Once the models were well-trained, we used the brain functional graphs of subjects in the A^+^ group to extract global A^+^-specific and A^-^-shared latent features through both encoders. We then performed representative similarity analysis (RSA)^133^ to identify the associations between these latent features and various characteristic variables. In RSA, we first utilized Euclidean distance to construct a subject similarity matrix based on z-score normalized characteristic variables including age, sex, MoCA, ADAS11, CDR, and others. Similarly, we constructed a subject similarity matrix based on the z-score normalized A^+^-specific and A^-^-shared latent features. The Mantel test (Spearmen correlation)^134^ was employed to compute the correlation between similarity matrices constructed from A^+^-specific latent features and those of characteristic features, as well as between the similarity matrices of A^-^-shared latent features and characteristic features. To further validate the uniqueness of the characteristic profiles between A^+^-specific and A^-^-shared features, we estimated associations between characteristic variables and latent features obtained from ablation models without the contrastive module, following the same steps, serving as a control study. To assess differences in significant correlations, Wilcoxon signed-rank tests were conducted, comparing Spearman correlation coefficients in the Mantel test between characteristic variables and target-specific, background-shared, and entangled latent features, across ten repeated runs of 5-fold cross-validated models. Upon detecting significant group-level differences, pairwise differences in correlations to characteristic variables across target-specific, background-shared, and entangled latent features were further identified using Dunn’s post-hoc test. All p values obtained from Dunn’s post-hoc test were FDR corrected across different characteristic variables.

### Transcriptomic and genomic analyses

#### Allen Human Brain Atlas dataset (AHBA)

The AHBA dataset offers high-resolution gene expression (RNA) data across 3702 brain spatially distinct tissue samples^14^. Samples were collected from six postmortem brains (age = 42.50 ± 13.38 years; male/female = 5/1), including T1 MRI data with MNI sample coordinates. The dataset was processed using the abagen package (https://abagen.readthedocs.io/en/stable/index.html) built in python^135^. Initially, microarray probes were matched to a valid Entrez ID based on the Entrez Gene database at the National Center for Biotechnology Information (NCBI)^136^. Probes with expression intensity below background noise^137^ in >= 50.00% of samples across donors were discarded, yielding 31569 probes. For genes indexed by multiple probes, the probe with the most consistent pattern of regional variation across donors was selected^138^. MNI coordinates were collected from Advanced Normalization Tools (ANTs; https://github.com/chrisfilo/alleninf), and samples were assigned to 100 brain regions defined in the Schaefer atlas if their MNI coordinates were within 2 mm of a given parcel. Tissue samples not assigned to a brain region in the atlas were discarded. Inter-subject variation was addressed by z-scoring tissue sample expression values across genes independently for each donor. Samples assigned to the same brain region were averaged separately for each donor and then across donors, resulting in a regional expression matrix (100 ROIs × 15633 genes).

#### Identifying genes linked to disentangled Aβ and p-tau variances

We used PLS regression to investigate genes associated with disentangled Aβ and p-tau variances captured by corresponding brain signatures (Figure S1 (D)). As a multivariate statistical method, PLS seeks to maximize the covariance between two variable sets and is advantageous in handling multicollinearity^139^. For each brain signature, one PLS model was calculated. In line with previous studies^19,49^, the first component of each PLS model, correlating with the brain signature, explained the largest variance in transcriptomic expression information (variance explained fraction for Aβ-associated A^+^-specific and A^-^-shared signatures were 64% and 49%; for p-tau-associated A^+^-specific and A^-^-shared signatures were 59% and 77%). The statistical significance of the PLS component was calculated using a permutation test by randomly permuting the expressed regions of genes 1000 times^49^. Additionally, a spin permutation test^140^ based on 10000 brain spatial spherical rotations was implemented to confirm its significance (Predicting Aβ, A^+^-specific signature: p_permute_ = 0.009, p_spin_ = 0.0001 and A^-^-shared signature: p_permute_ = 0.049, p_spin_ = 0.0001; Predicting p-tau, A^+^-specific signature: p_permute_ = 0.036, p_spin_ = 0.0001 and A^-^-shared signature: p_permute_ = 0.001, p_spin_ = 0.0001). To assess the significance of each gene’s contribution to the first PLS component, we bootstrapped the brain regions 2000 times (two-tailed test, the null hypothesis is that the weight of this gene equal to zero) and corrected the significance of the weight across all genes (15633) using FDR. Only significantly weighted genes (p_FDR_ ≤ 0.05) were selected for further analysis. To fairly compare the weight of each gene, the loading weights of PLS models were divided by the standard deviation of the bootstrap distribution to calculate Z scores. We ranked the genes based on Z scores to determine their contribution to the PLS model. The similarity of the significantly weighted gene lists across all PLS models was assessed using rank biased overlap (RBO), an indefinite rank similarity measurement designed to handle non-conjoint ranking lists^48^.

#### Enrichment analysis of identified gene lists

For the enrichment analysis of identified gene lists, we used Metascape (https://metascape.org/gp/index.html#/main/step1), a web-based portal designed to provide functional enrichment, interactome analysis, gene annotation over 40 independent knowledgebases^39^. In our study, four well-established gene annotation databases: ontology (GO) biological processes, Kyoto Encyclopedia of Genes and Genomes (KEGG), Reactome, and Wiki pathways were included. Utilizing this portal, we performed GSEA for all gene lists separately to obtain the top biological functional pathways linked to Aβ and p-tau-related brain regional signatures. Subsequently, we employed RBO to measure the similarity of enriched pathways between the identified gene list from the PLS model and gene list revealed from the established GWAS of AD^47^, to elucidate the consistency between our finding and reported finding. Additionally, a multi-gene-list meta-analysis was performed to obtain shared enriched pathways involving those gene lists.

#### Assigning Aβ and p-tau correlated gene lists to cell types

Given the association between cellular diversity and complex interactions in the brain and AD development, we aimed to explore the genes associated with CSF neuropathological protein expression and canonical cell classes of the central nervous system. Gene lists involving metabolism of seven canonical cell classes: microglia, endothelial cells, oligodendrocyte precursors, oligodendrocytes, astrocytes, excitatory, and inhibitory neurons were obtained from previous gene clustering results initialized in 58 cell classes^51^. According to this prior knowledge, PLS-identified gene lists were assigned to the seven canonical cell types. The significance of overlapping counts between gene lists identified by PLS and seven canonical cell-related gene lists was determined using a permutation test of PLS regression conducted 1000 times. Subsequently, a multi-gene-list meta-analysis was implemented to detect the enriched molecular pathways specific to different cell types linked to CSF Aβ and p-tau expression.

## Data availability

The data supporting the results in this study are available within the paper and its Supplementary Information. The PREVENT-AD dataset is publicly available (https://openpreventad.loris.ca/). The ADNI dataset is publicly available (https://adni.loni.usc.edu/).

## Code availability

All analyses were implemented in Python 3.9. The statistical analyses were implemented using scipy 1.9.3(https://docs.scipy.org/doc/scipy-1.9.3/). The deep graph model was built using Pytorch (2.0.1+cu118, https://pytorch.org/), PyG (2.3.1, https://www.pyg.org/). Resting-state functional MRI data were processed with fMRIPrep 20.2.3 (https://hub.docker.com/r/nipreps/fmriprep/tags). Internal operations of fMRIPrep 20.2.3 use the following software: Advanced Normalization Tools 2.3.3, Nipype 1.6.1, FSL 5.0.9, FreeSurfer 6.0.1, AFNI 20160207. The code will be made available upon the publication of this paper.

## Acknowledgements

This work was supported by NIH grant nos. R21AG080425, R01MH129694, R21MH130956, Alzheimer’s Association Grant (AARG-22-972541), and Lehigh University FIG (FIGAWD35), CORE, and Accelerator grants. Portions of this research were conducted on Lehigh University’s Research Computing infrastructure partially supported by NSF Award 2019035. G.A.F. was also supported by philanthropic funding and NIH grant nos. R01MH132784 and R01MH125886, and grants from the One Mind - Baszucki Brain Research Fund, the SEAL Future Foundation, and the Brain and Behavior Research Foundation. This study was supported by the Fondo de Investigaciones Sanitario, Carlos III Health Institute (INT21/00073, PI20/01473 and PI23/01786 to J.F.) and the Centro de Investigación Biomédica en Red sobre Enfermedades Neurodegenerativas Program 1, partly jointly funded by Fondo Europeo de Desarrollo Regional, Unión Europea, Una Manera de Hacer Europa. This work was also supported by the National Institutes of Health grants (R01 AG056850; R21 AG056974, R01 AG061566, R01 AG081394 and R61AG066543 to J.F.), the Department de Salut de la Generalitat de Catalunya, Pla Estratègic de Recerca I Innovació en Salut (SLT006/17/00119 to J.F.). It was also supported by Fundación Tatiana Pérez de Guzmán el Bueno (IIBSP-DOW-2020-151 o J.F.) and Horizon 2020– Research and Innovation Framework Programme from the European Union (H2020-SC1-BHC-2018- 2020 to J.F.).

Data collection and sharing for this project was funded by the Alzheimer’s Disease Neuroimaging Initiative (ADNI) (National Institutes of Health Grant U01 AG024904) and DOD ADNI (Department of Defense award number W81XWH-12-2-0012). ADNI is funded by the National Institute on Aging, the National Institute of Biomedical Imaging and Bioengineering, and through generous contributions from the following: AbbVie, Alzheimer’s Association; Alzheimer’s Drug Discovery Foundation; Araclon Biotech; BioClinica, Inc.; Biogen; Bristol-Myers Squibb Company; CereSpir, Inc.; Cogstate; Eisai Inc.; Elan Pharmaceuticals, Inc.; Eli Lilly and Company; EuroImmun; F. Hoffmann-La Roche Ltd and its affiliated company Genentech, Inc.; Fujirebio; GE Healthcare; IXICO Ltd.; Janssen Alzheimer Immunotherapy Research & Development, LLC.; Johnson & Johnson Pharmaceutical Research & Development LLC.; Lumosity; Lundbeck; Merck & Co., Inc.; Meso Scale Diagnostics, LLC.; NeuroRx Research; Neurotrack Technologies; Novartis Pharmaceuticals Corporation; Pfizer Inc.; Piramal Imaging; Servier; Takeda Pharmaceutical Company; and Transition Therapeutics. The Canadian Institutes of Health Research is providing funds to support ADNI clinical sites in Canada. Private sector contributions are facilitated by the Foundation for the National Institutes of Health (www.fnih.org). The grantee organization is the Northern California Institute for Research and Education, and the study is coordinated by the Alzheimer’s Therapeutic Research Institute at the University of Southern California. ADNI data are disseminated by the Laboratory for Neuro Imaging at the University of Southern California.

