## Supplementary Material for "Elucidating the neuropathological and molecular heterogeneity of amyloid-β and tau in Alzheimer’s disease through machine learning and transcriptomic integration"

### Supplementary Materials

**Figure S1. Workflow of this study. (A)** Brain functional graph construction. We calculated Pearson correlation coefficients across all 100 brain regions as functional connectivity defined by Schaefer parcellation. Subsequently, the raw FC matrix was utilized as the node feature, while the binarized FC matrix served as the adjacency matrix of brain functional graph. **(B)** Model training. We input both target dataset and all datasets containing both target and background datasets into two separate encoders with identical structures. The model in pretraining phase with contrast loss to disentangle the latent features encoded from two encoders. Then we finetune the model to predict the true scores from target dataset and background dataset.

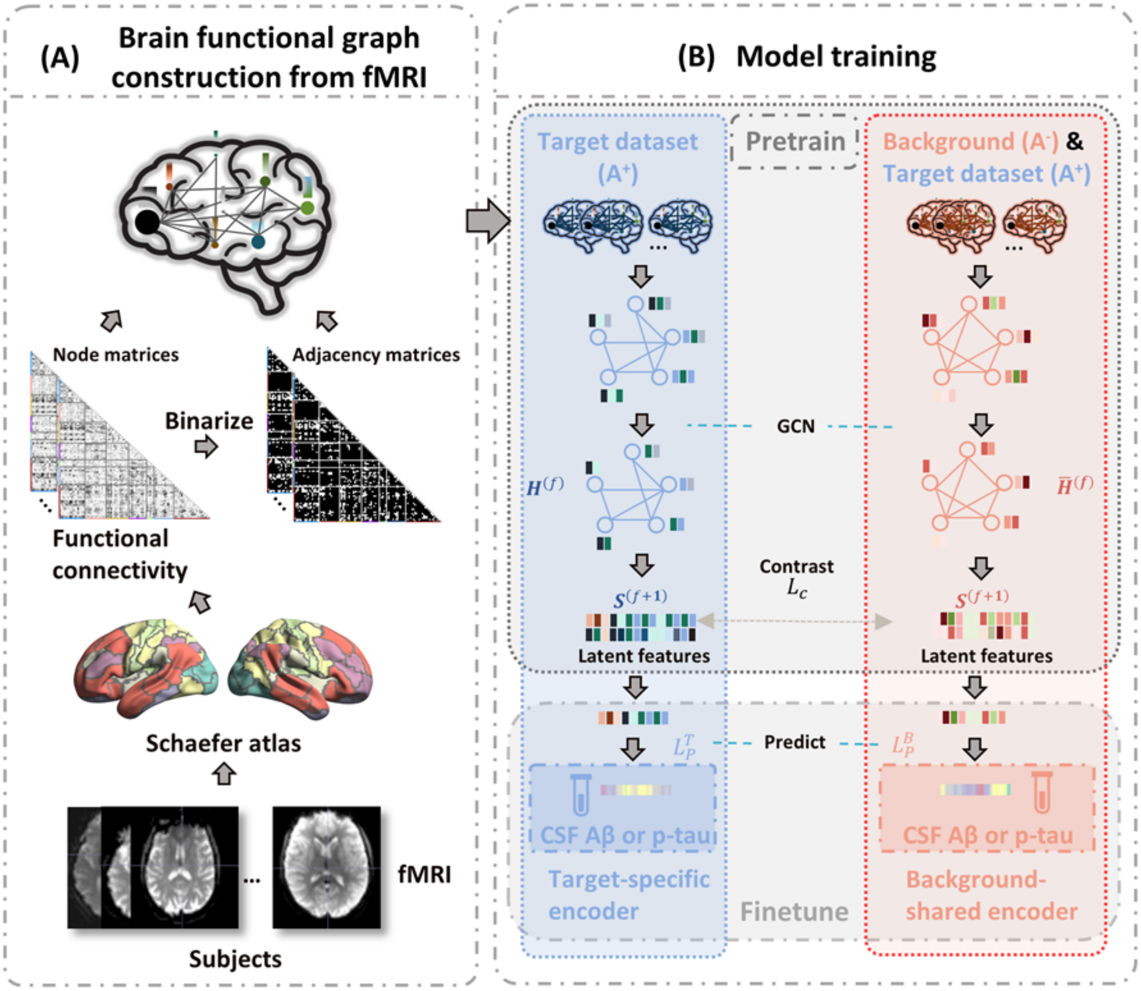

**Figure S2. Post-hoc exploration of this study. (A)** Representative similarity analysis. We input the target dataset ($A^+$) into the two encoders of well-trained models to obtain target-specific and background-shared latent features. Similarity matrices of z-scored characteristic variables, target-specific and background-shared latent features were calculated using Euclidean distance. The associations between these similarity matrices were assessed using the Mantel test (Spearman). **(B)** Transcriptomic association analyses. First, we employed a partial least squares (PLS) model to establish associations between whole brain gene expression patterns and brain signatures predictive for CSF $A\beta_{1\text{–}42}$ and p-$tau_{181p}$ from different encoders. Second, the significance of PLS first component was confirmed using simple permutation test and spatial permutation ('spin') test. Bootstrapping was then conducted to identify the gene set significantly contributing to the PLS first component and those genes were visualized in ascending order. Third, gene enrichment analysis summarized the aggregated ontological pathways of the obtained gene set.

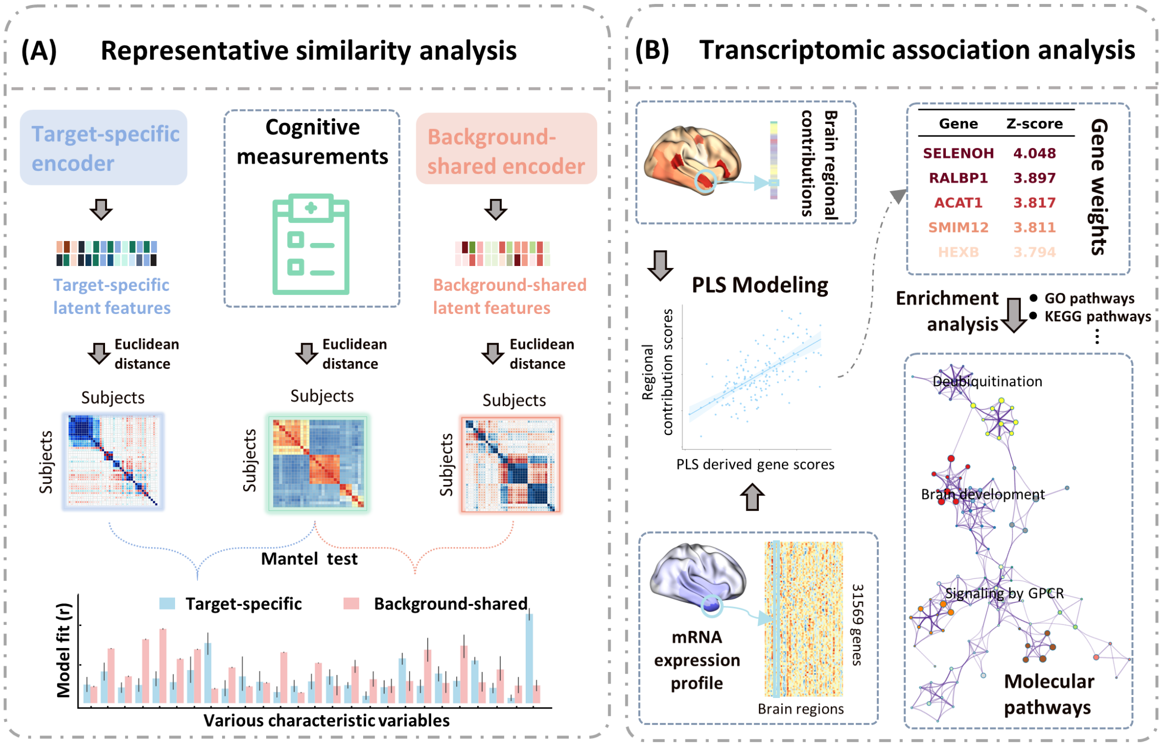

**Figure S3. Characteristic profiles of CSF Aβ and p-tau linked brain latent features of $A^+$ subjects in cognitive unimpaired (CU), mild cognitive impairment (MCI) and dementia subgroups.** Representative similarity analysis was used to estimate the associations between characteristic variables and latent features, obtained from 10 × 5-fold cross-validated models in predicting CSF Aβ and p-tau separately. Data expressed as mean ± s.d. Fisher's combined probability test confirmed the significance of the correlation between each characteristic variable and latent features disentangled from $A^+$-specific, $A^-$-shared encoders, and ablation models without contrasting module in 10 trials. Kruskal-Wallis one-way ANOVA was utilized to detect group differences in correlations involving latent features disentangled from $A^+$-specific, $A^-$-shared encoders, and ablation models without contrasting module. Dunn's test was then employed to detect pairwise differences in correlation. False discovery rate (FDR) correction of statistic comparison across all correlations involving multiple characteristic variables and latent features was applied. Significance levels for pairwise differences were denoted by *($p \leq 0.05$), **($p \leq 0.01$), ***($p \leq 0.001$), and ****($p \leq 0.0001$). **(A, B)** Associated characteristic profiles to $A^+$-specific, $A^-$-shared and entangled brain graphs latent features of subjects in $A^+$ CU group, guided by the CSF **(A)** Aβ and **(B)** p-tau expressions. **(C, D)** Associated characteristic profiles to $A^+$-specific, $A^-$-shared and entangled brain graphs latent features of subjects in $A^+$ MCI group, guided by the CSF **(C)** Aβ **(D)** p-tau expressions. **(E, F)** Associated characteristic profiles to $A^+$-specific, $A^-$-shared and entangled brain graphs latent features of subjects in $A^+$ dementia group, guided by the CSF **(E)** Aβ **(F)** p-tau expressions.

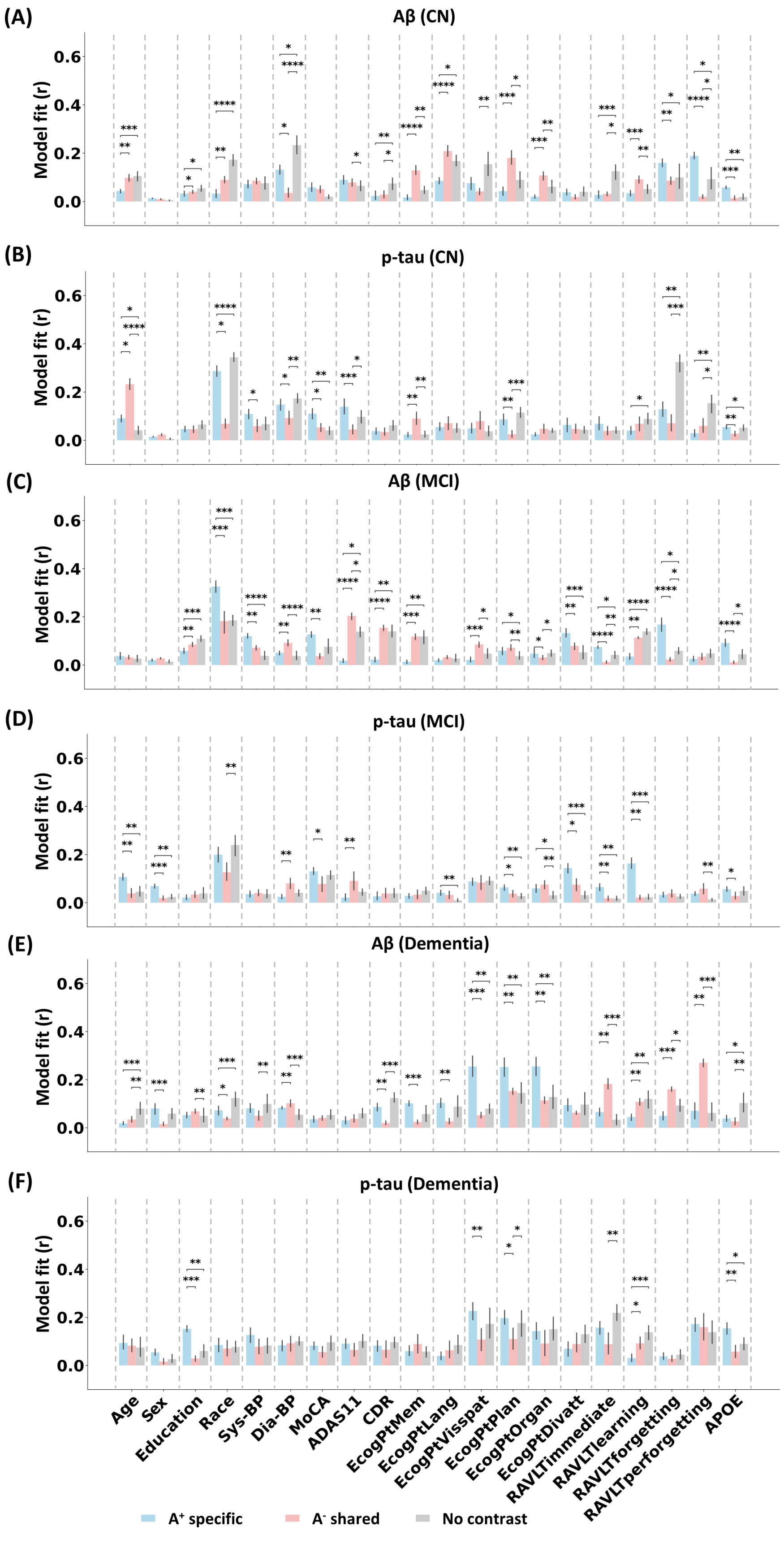

(A)
Aβ (CN)
(B)
p-tau (CN)
(C)
Aβ (MCI)
(D)
p-tau (MCI)
(E)
Aβ (Dementia)
(F)
p-tau (Dementia)
Model fit (r)
Age
Sex
Education
Race
Sys-BP
Dia-BP
MoCA
ADAS11
CDR
EcogPtMem
EcogPtLang
EcogPtVisspat
EcogPtPlan
EcogPtOrgan
EcogPtDivatt
RAVLTimmediate
RAVLTlearning
RAVLTforgetting
RAVLTperforgetting
APOE
A+ specific
A- shared
No contrast

**Figure S4. Characteristic profiles of CSF Aβ and p-tau linked brain latent features in $A^+T^-$ subgroup.** Representative similarity analysis was used to estimate the associations between characteristic variables and latent features, obtained from 10 × 5-fold cross-validated models in predicting CSF Aβ and p-tau separately. Data expressed as mean ± s.d. Fisher's combined probability test confirmed the significance of the correlation between each characteristic variable and latent features disentangled from $A^+$-specific, $A^-$-shared encoders, and ablation models without contrasting module in 10 trials. Kruskal-Wallis one-way ANOVA was utilized to detect group differences in correlations involving latent features disentangled from $A^+$-specific, $A^-$-shared encoders, and ablation models without contrasting module. Dunn's test was then employed to detect pairwise differences in correlation. False discovery rate (FDR) correction of statistic comparison across all correlations involving multiple characteristic variables and latent features was applied. Significance levels for pairwise differences were denoted by *($p \leq 0.05$), **($p \leq 0.01$), ***($p \leq 0.001$), and ****($p \leq 0.0001$). **(A)** Associated characteristic profiles to $A^+$-specific, $A^-$-shared and entangled brain graphs latent features, guided by prediction of Aβ expressions. **(B)** Associated characteristic profiles to $A^+$-specific, $A^-$-shared and entangled brain graphs latent features, guided by prediction of p-tau expressions.

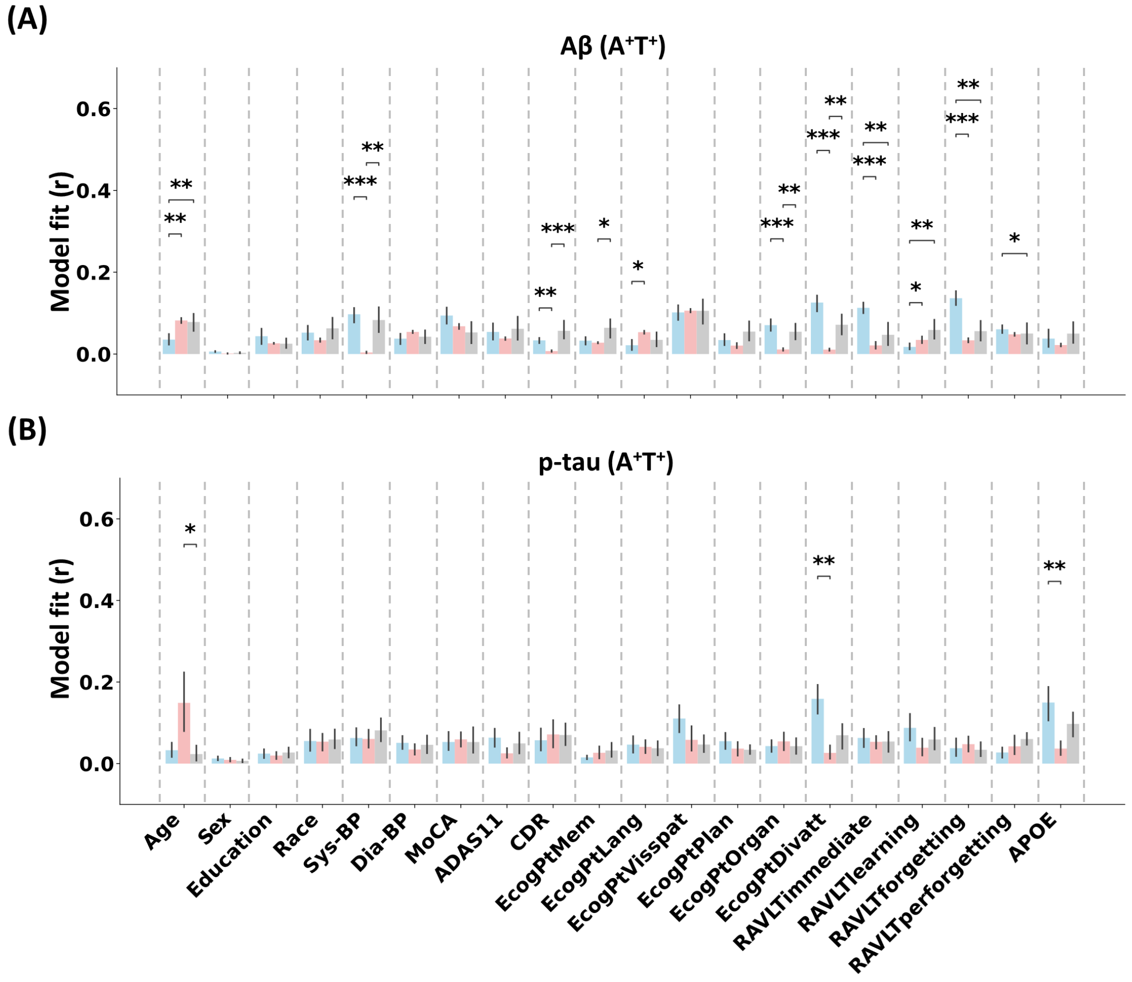

**Figure S5. The network plot of top 20 enriched functional terms of the significantly correlated genes for each brain signature. (A)** Network plot of the top 20 enriched terms of the significantly correlated genes of $A^+$-specific brain signature in predicting Aβ. **(B)** Network plot of the top 20 enriched terms of the significantly correlated genes of $A^-$-shared brain signature in predicting Aβ. **(C)** Network plot of the top 20 enriched terms of the significantly correlated genes of $A^+$-specific brain signature in predicting p-tau. **(D)** Network plot of the top 20 enriched terms of the significantly correlated genes of $A^-$-shared brain signature in predicting p-tau.

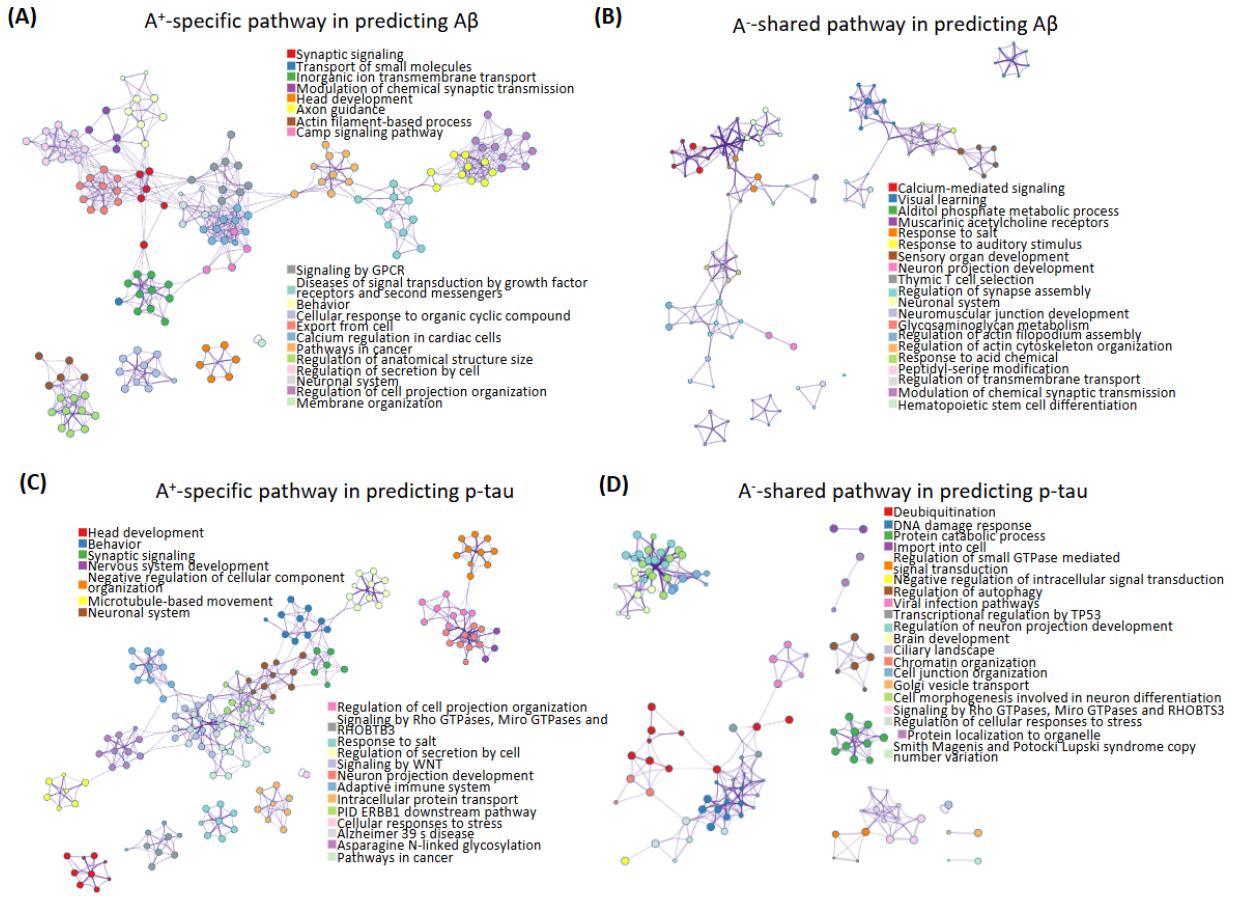

**Figure S6. Associations across gene lists linked to different signatures and genome-wide association (GWAS) of AD. (A)** Heatmap of similarity across ranked gene lists. This heatmap displays the similarity across various gene lists linked to CSF Aβ linked and p-tau linked variances, and those identified in previous GWAS. The similarity distance was evaluated by rank biased overlap (RBO). **(B)** Chord plot of gene lists. **(C)** Heatmap and **(D)** network plot of the top 20 overlapping enriched terms across the gene lists linked to CSF Aβ and p-tau variances and those identified in previous GWAS. Overall, the gene lists exhibit a high correlation with CSF Aβ and p-tau linked $A^{+}$-specific features and those two gene lists show some overlapping to the gene list of established GWAS.

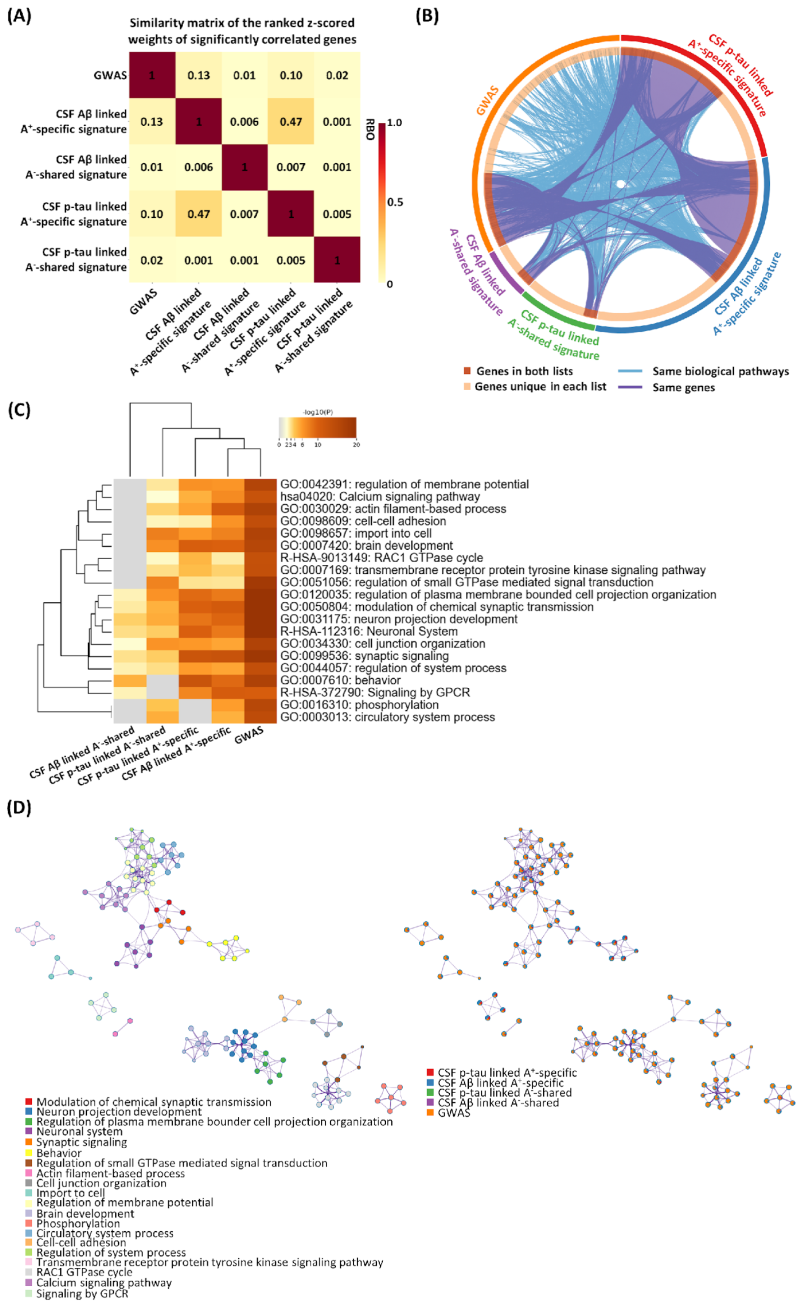

**Figure S7. Comparison of performance between our model and other typical machine learning models.** Bars expressed as mean ± std of $R^2$ from one repetition of 10-fold cross-validation. $R^2$ of our models were 0.15 ± 0.02 and 0.16 ± 0.02 for Aβ and p-tau prediction respectively. We then used Wilcoxon signed-rank test to compared the $R^2$ values of our model against those of the next best-performing method, affirming its significant superiority (Aβ prediction: Z = 2.15, p = 0.03; p-tau prediction: Z = 2.23, p = 0.02).

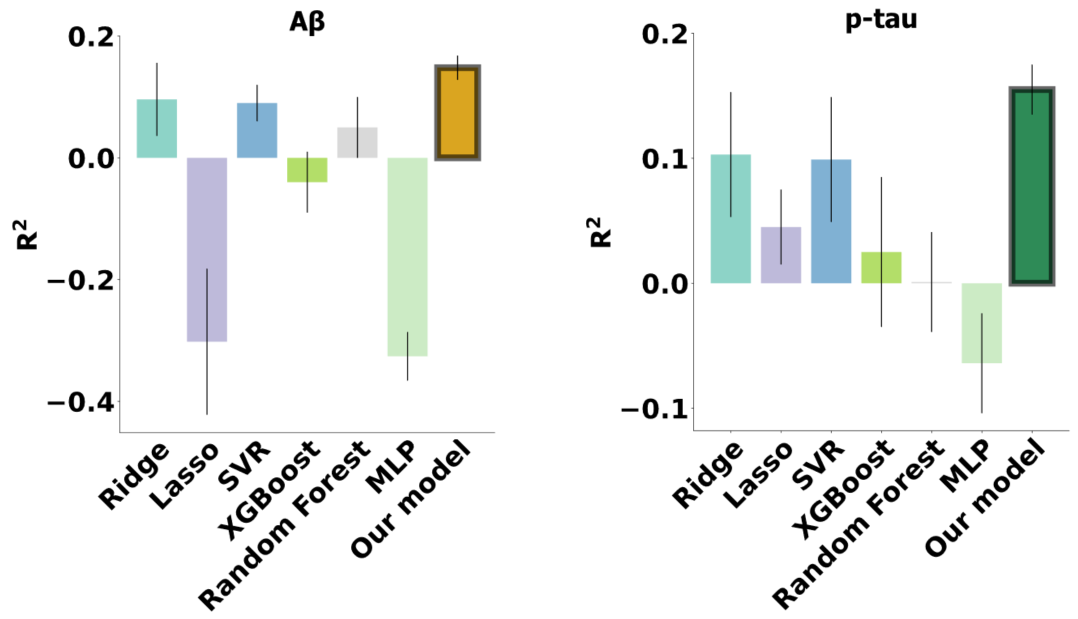

**Figure S8. Ablation study of our model.** We iteratively deleted one of the different modules in our model to verify the contribution of this module to the prediction performance of our model. Bars expressed as mean ± s.d. of evaluated score from one cross-validation trial (n = 10). The term "*MLP instead of dGCN*" indicates the utilization of the same model module with the substitution of MLP instead of a GCN. "*No contrast block*" signifies training our model without contrasting the background group from the target group ($A^+$) and "*No pretrain*" indicates training our model without the pretraining steps aimed at maximizing mutual information between original brain graph and the latent features of target group.

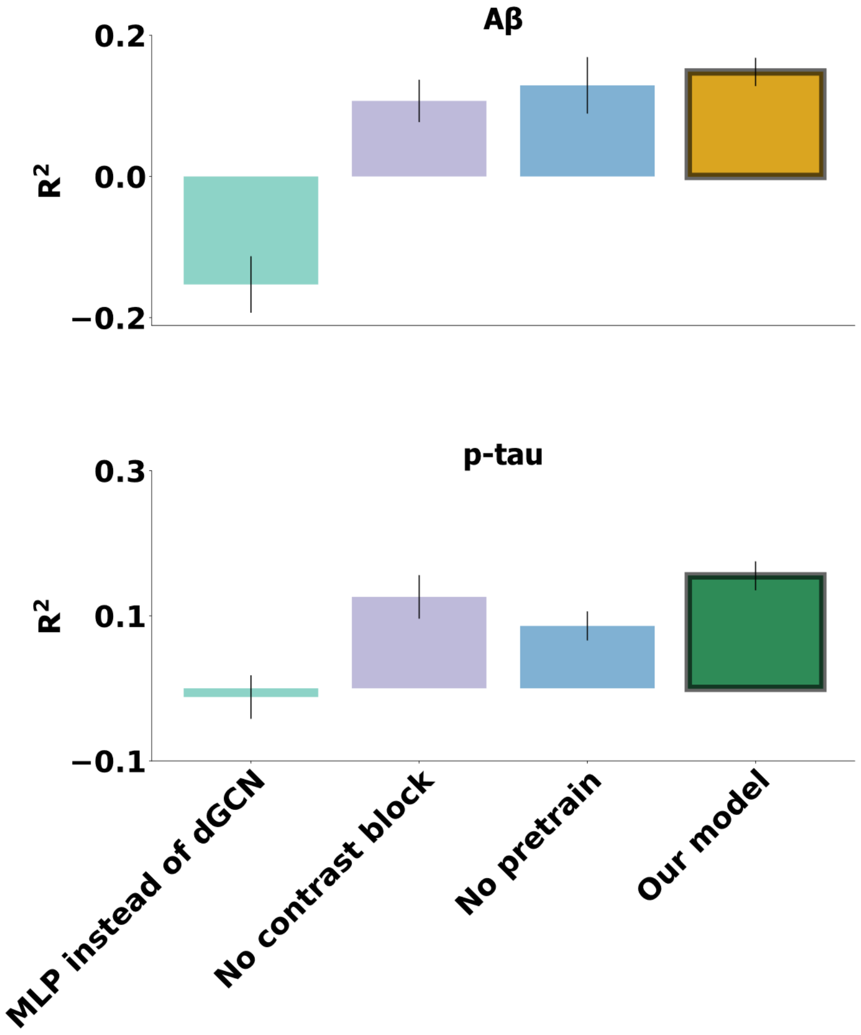

**Figure S9. Prediction performance when we matched the sex and age in A⁻.** We randomly selected the subgroup from A⁻ subjects with matched demographic information and retrained the models in 10 × 5-fold cross validation. **(A)** $R^2$ between predicted CSF Aβ and true CSF Aβ expression, obtained from our models training with matched demographic information or not. Wilcoxon signed-rank test was used to compared the $R^2$ values ($Z = -0.77$, $p = 0.45$). **(B)** The scatter plot between ensembled predicted CSF Aβ and true CSF Aβ expression, matching demographic information. **(C)** $R^2$ between predicted CSF p-tau and true CSF p-tau expression, obtained from our models training with matched demographic information or not. Wilcoxon signed-rank test was used to compared the $R^2$ values ($Z = -0.76$, $p = 0.46$). **(D)** The scatter plot between ensembled predicted CSF p-tau and true CSF p-tau expression, matching demographic information.

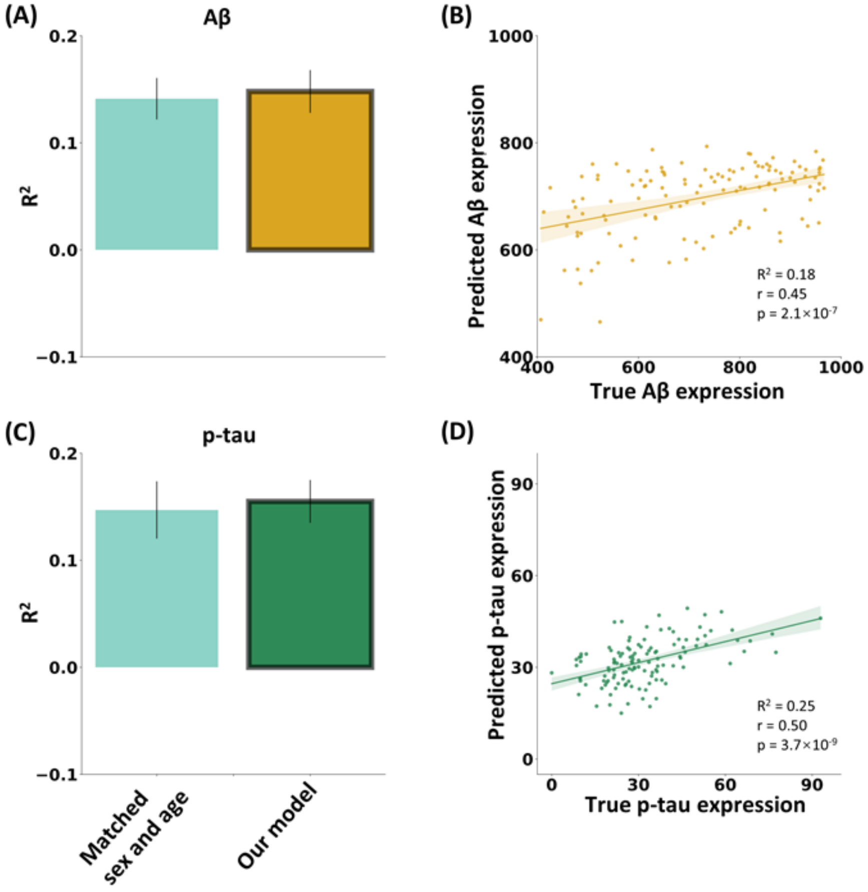

**Figure S10. Prediction performance evaluated by partial correlation between predicted and true scores when we controlled for the site labels.** We evaluated the partial correlation in 10 trials of 5-fold cross validation. No significant change in prediction performance was observed after controlling for site labels (Aβ prediction: Z = -0.08, p = 0.94; p-tau prediction: Z = -0.53, p = 0.60).

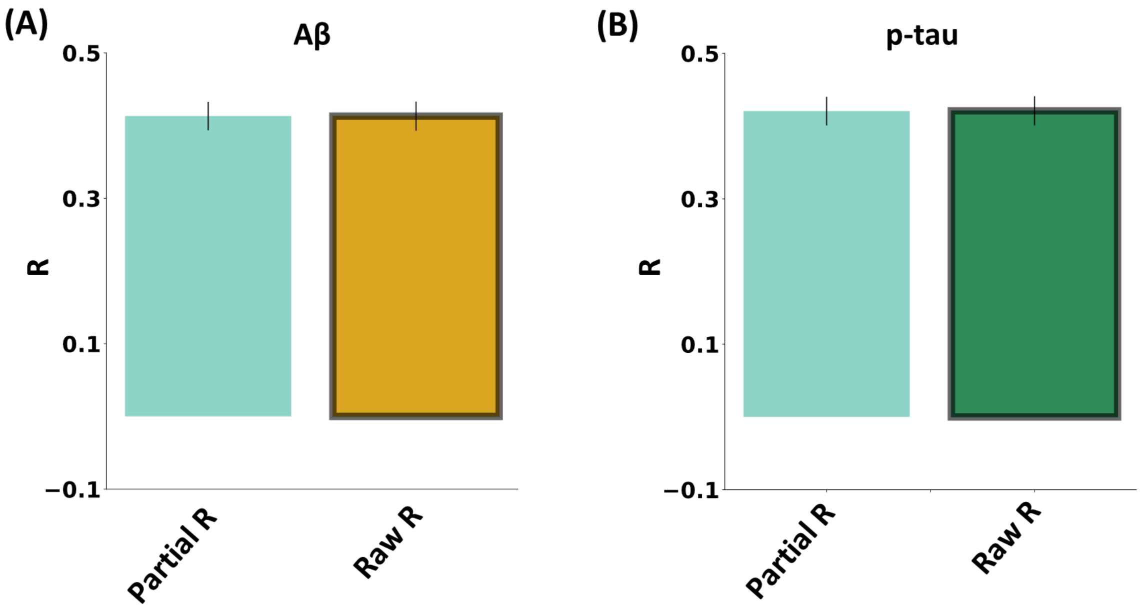

**Figure S11. Prediction performance for various AD neuropathological protein expression of combination of $A^*T^+$ and $A^+T^*$ groups across different models using 10 × 5-fold cross validation.** The target group ($A^*T^+$ and $A^+T^*$) was defined based on the CSF Aβ > 976.6 pg/ml or p-tau < 21.8 pg/ml. The panels are organized with the left side of the dashed line presenting results from well-established machine learning models, while the right side showcases findings from ablation studies on our models. Specifically, "*MLP instead of dGCN*" denotes the substitution of MLP instead of GCN within the same model module. "*No contrast block*" signifies training our model without contrasting the background group (all subjects) from the target group ($A^*T^+$ and $A^+T^*$), and "*no pretrain*" indicates training our model without the pretraining steps aimed at maximizing mutual information between brain graph embeddings and the latent features of the target group. **(A)** $R^2$ between predicted CSF Aβ and true CSF Aβ expression, obtained from different models. **(B)** The scatter plot between ensembled predicted CSF Aβ and true CSF Aβ expression, using our models from 10 × 5-fold cross validations. **(C)** $R^2$ between predicted CSF p-tau and true CSF p-tau expression, obtained from different models. **(D)** The scatter plot between ensembled predicted CSF p-tau and true CSF p-tau expression, using our models from 10 × 5-fold cross validations.

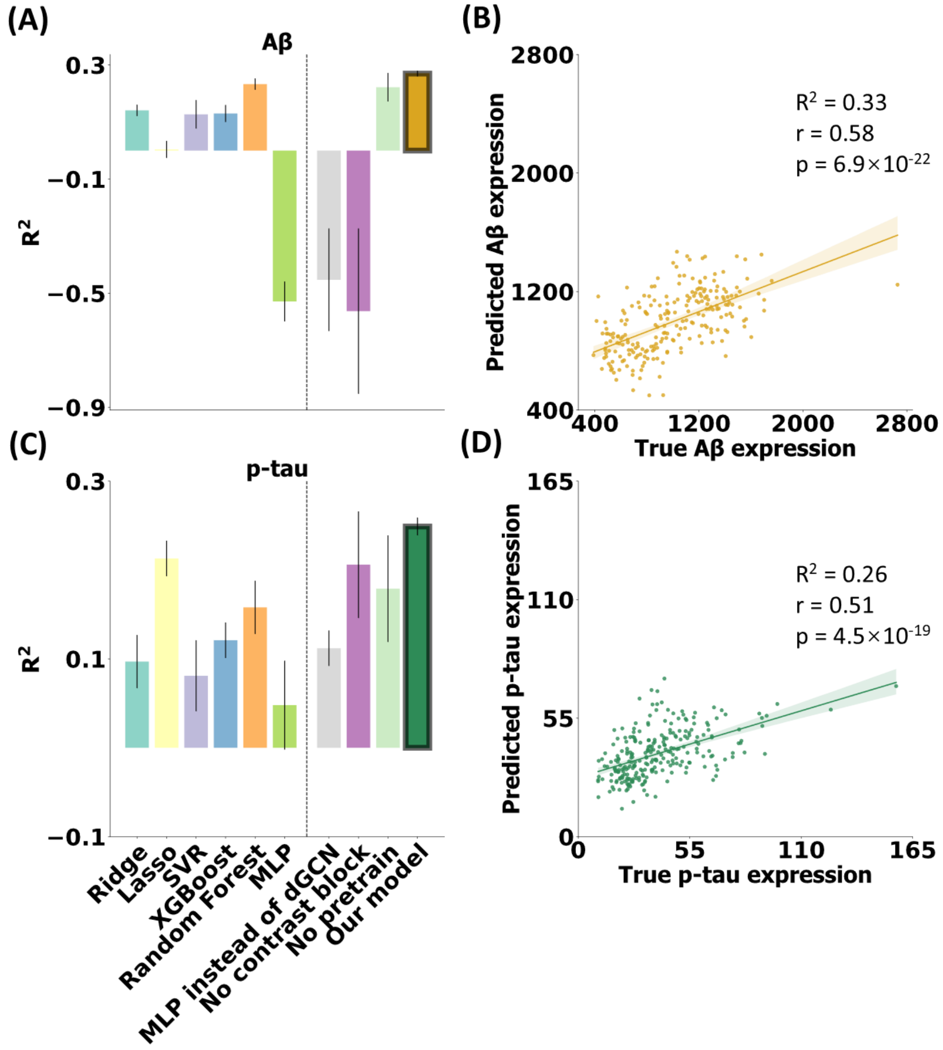

**Figure S12. Prediction performance for various AD neuropathological protein expressions across different models using 10 × 5-fold cross-validation underlying dementia&MCI/CU framework.** The target group was defined as dementia and MCI and the background group was defined as CU. The term "MLP instead of dGCN" indicates the utilization of the same model module with the substitution of MLP instead of a GCN. "No contrast block" signifies training our model without contrasting the background group from the target group (dementia&MCI) and "no pretrain" indicates training our model without the pretraining steps aimed at maximizing mutual information between brain graph and the latent features of the target group. **(A)** $R^2$ between predicted CSF Aβ and true CSF Aβ expression, obtained from different models. **(B)** The scatter plot between ensembled predicted CSF Aβ and true CSF Aβ expression, using our models from 10 × 5-fold cross validations. **(C)** Pearson correlation between predicted CSF Aβ and true CSF Aβ expression obtained from different models. **(D)** The scatter plot between ensembled predicted CSF p-tau and true CSF p-tau expression, using our models from 10 × 5-fold cross validations.

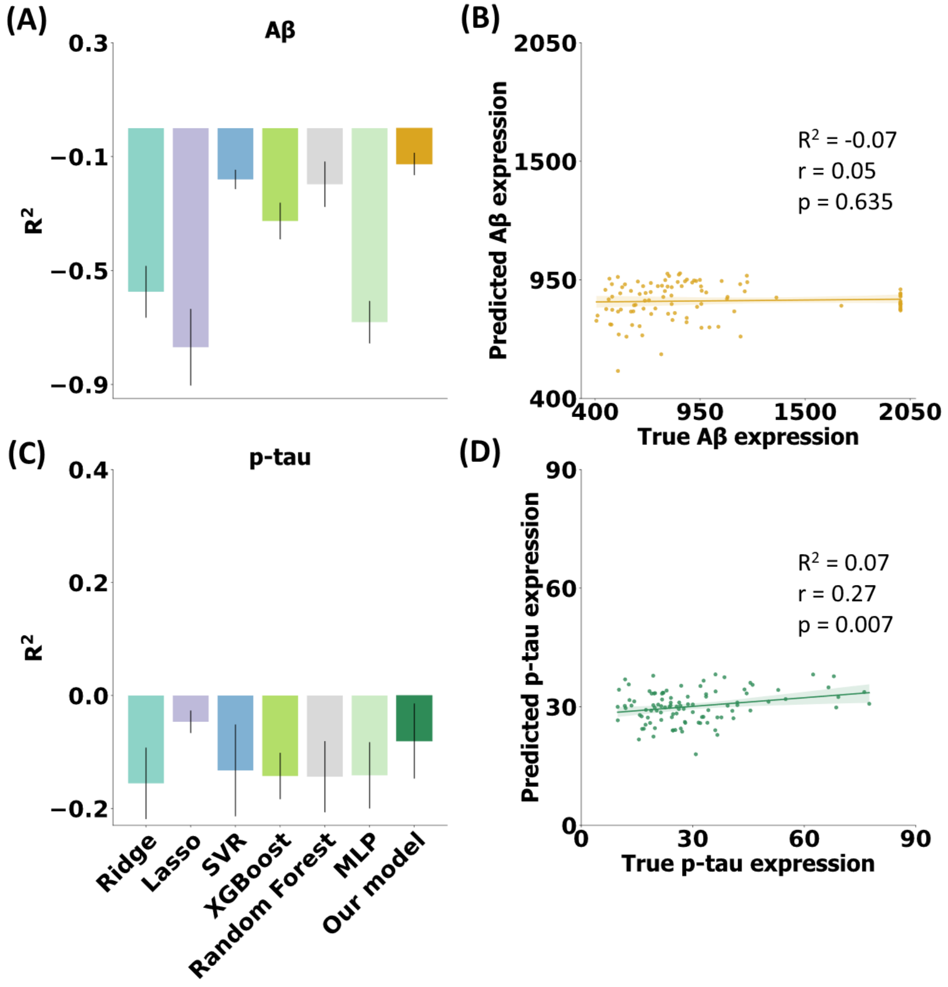

**Figure S13. Contrastive effect in predicting various neuropathological proteins. (A)** The comparison of distance of latent features within A$^+$ group and between A$^+$/A$^-$ groups. We first used Jensen-Shannon divergence to measure the distance of latent features between A$^+$ subjects and A$^-$ subjects, encoding from our cGCN model. We then measured the distance of latent features between A$^+$ subjects and A$^-$ subjects encoding from the ablation GCN without contrasting. The two-sample t-test was used to detect the difference of Jensen-Shannon divergences within A$^+$ group and between A$^+$ and A$^-$ groups. **(B)** The comparison of graph properties of features between A$^+$ and A$^-$ groups. We calculated average clustering coefficient, modularity and path length of the graphs constructed from raw FC and latent node features predictive for Aβ and p-tau of A$^+$ and A$^-$ groups separately. The two-sample t-test was used to compare the difference of these graph coefficients between A$^+$ and A$^-$ groups. *($p \leq 0.05$), **($p \leq 0.01$), ***($p \leq 0.001$), and ****($p \leq 0.0001$), NS (no significance). Lowercase d denoted Cohen's d, indicating the standardized difference between means from two compared groups.

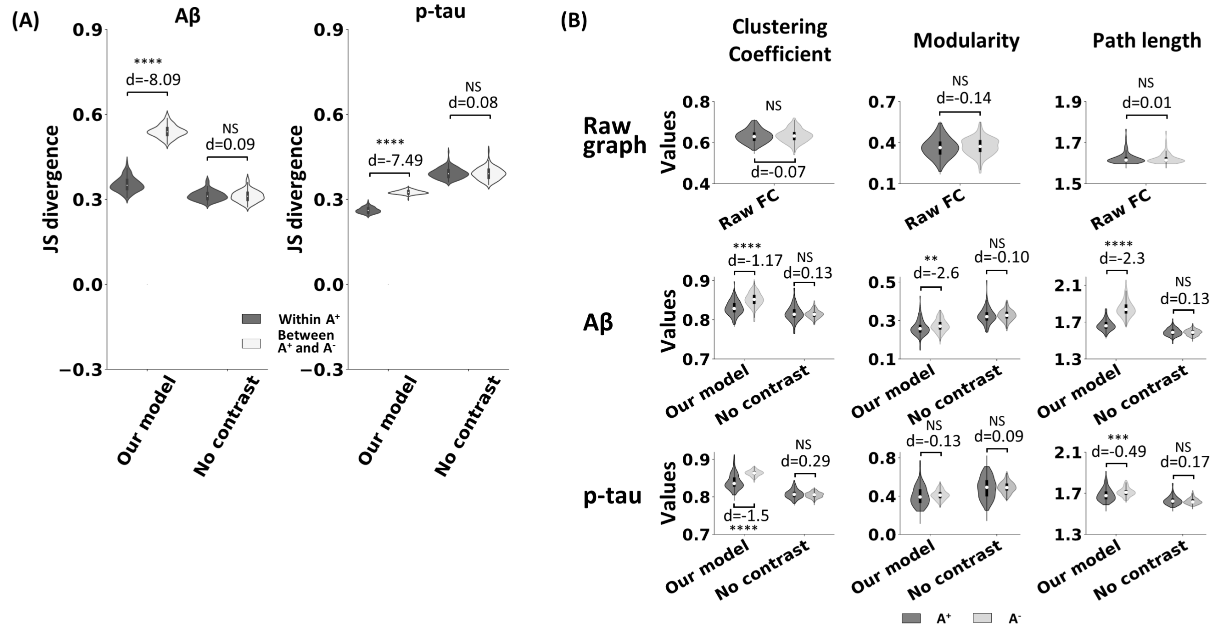

**Figure S14. Prediction using well-trained A⁺-specific encoder for A⁻ subjects. (A)** The scatter plot between predicted Aβ averaged from ten trials of 5-fold cross validations and true Aβ expression. **(B)** The scatter plot between predicted p-tau averaged from ten trials of 5-fold cross validations and true p-tau expression.

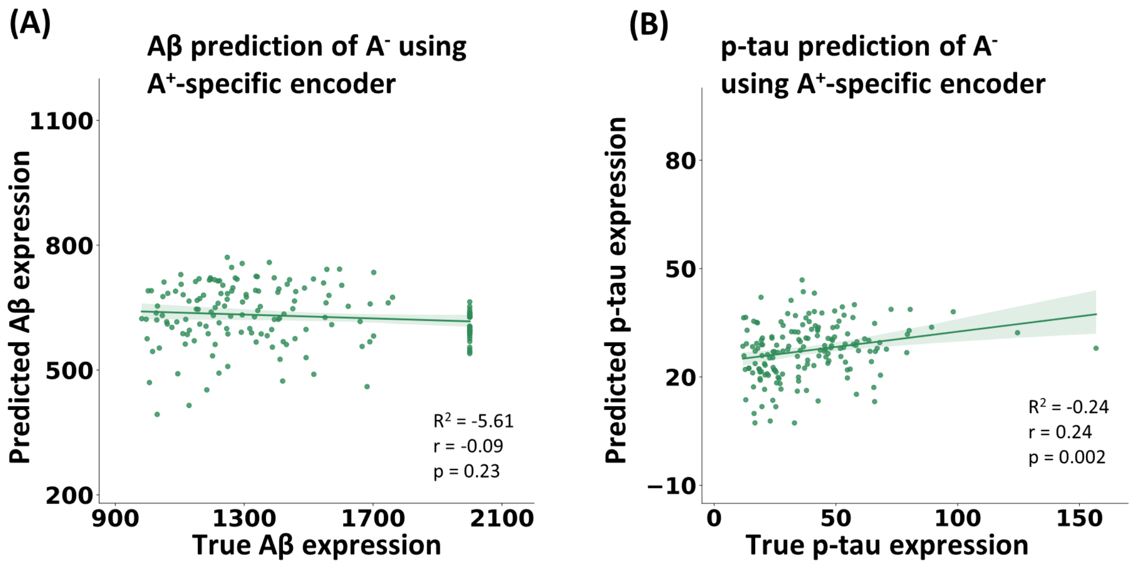

**Figure S15. Counts of FCs significantly correlated to neuropathological proteins ($p_{FDR} < 0.05$) differential in target and background groups.** We measured the Pearson correlation between node features (FCs) and predicting neuropathological proteins in target and background groups defined using different frameworks. We then defined specific correlated FC sets in target group as the FCs which uniquely correlated to neuropathological proteins in target group or correlated reversely between target and background group. Finally, we showed the counts of specific correlated FC sets in target group.

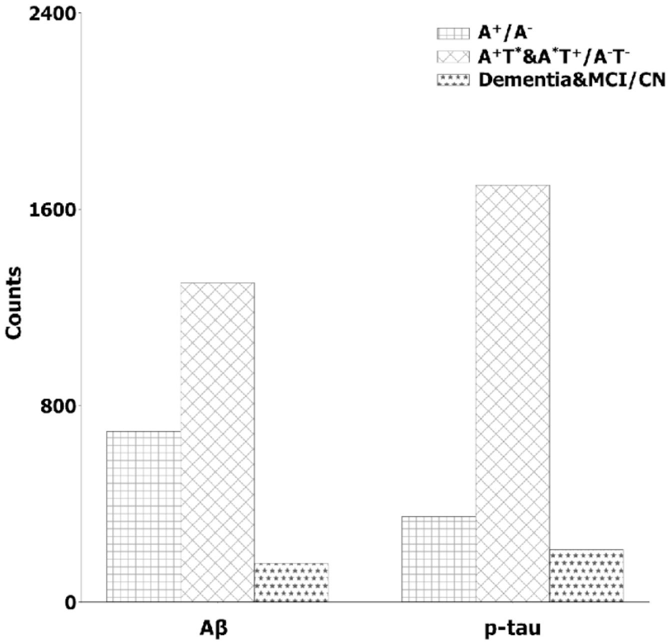

**Table S1. Summary of demographic information for subjects with CSF Aβ expression level as a cutoff to define target and background groups in this study.** The target group was defined as $A^+$, based on CSF $A\beta_{1\text{-}42} < 976.6$ pg/ml.

| ADNI & PREVENT-AD | $A^*$ contrast strategy | | | | | |
|---|---|---|---|---|---|---|
| | $A^+$ | | $A^-$ | | Statistic values | |
| | n | % | n | % | χ2 | p |
| Sex | | | | | | |
| Male | 62 | 48 | 106 | 64 | 6.8 | 0.009 |
| Female | 67 | 52 | 60 | 36 | | |
| Race | | | | | | |
| Asian | 2 | 2 | 3 | 2 | 5.3 | 0.15 |
| Black | 7 | 5 | 3 | 2 | | |
| White | 115 | 89 | 158 | 95 | | |
| Unknown | 5 | 4 | 2 | 1 | | |
| | mean | std | mean | std | t | p |
| Age (year) | 70.4 | ± 7.1 | 66.3 | ± 7.6 | 5.1 | $7\times10^{-7}$ |
| Education year | 15.4 | ± 2.6 | 15.9 | ± 2.9 | -1.2 | 0.21 |

**Table S2. Summary of demographic information for subjects in ADNI dataset.**

| Demographic variables | $A^+$ | | $A^-$ | | Statistic values | |
|---|---|---|---|---|---|---|
| | n | % | n | % | χ2 | p |
| Sex | | | | | | |
| Male | 50 | 48 | 28 | 64 | 0.001 | 0.98 |
| Female | 52 | 52 | 31 | 36 | | |
| Race | | | | | | |
| Asian | 2 | 0 | 3 | 2 | 2.6 | 0.46 |
| Black | 7 | 0 | 3 | 2 | | |
| White | 88 | 0 | 58 | 95 | | |
| Unknown | 5 | 0 | 1 | 1 | | |
| | mean | std | mean | std | t | p |
| Age (year) | 72.5 | ± 6.7 | 71.0 | ± 6.9 | 1..4 | 0.16 |
| Education year | 15.9 | ± 2.6 | 16.4 | ± 2.6 | -1.4 | 0.17 |

**Table S3. Summary of demographic information for subjects in PREVENT-AD dataset.**

| Demographic variables | A+ | | A- | | Statistic values | |
|---|---|---|---|---|---|---|
| | n | % | n | % | $\chi 2$ | p |
| Sex | | | | | | |
| Male | 13 | 48 | 29 | 29 | 2.8 | 0.09 |
| Female | 14 | 52 | 72 | 71 | | |
| Race | | | | | | |
| Asian | 0 | 0 | 0 | 0 | 0 | 1 |
| Black | 0 | 0 | 0 | 0 | | |
| White | 27 | 100 | 100 | 99 | | |
| Unknown | 0 | 0 | 1 | 1 | | |
| | mean | std | mean | std | t | p |
| Age (year) | 62.0 | ± 4.6 | 63.2 | ± 5.3 | -1.0 | 0.30 |
| Education year | 13.7 | ± 2.2 | 15.6 | ± 3.0 | -2.9 | 0.004 |

**Table S4. Summary of demographic information for subjects with CSF Aβ and p-tau expression level as a cutoff to define target and background groups in this study.** The target group was defined as $A^{+}T^{*}$ and $A^{*}T^{+}$, based on CSF $A\beta_{1\text{-}42}$ < 976.6 pg/ml ($A^{+}T^{*}$) or CSF p-tau > 21.8 pg/ml ($A^{*}T^{+}$).

| **ADNI & PREVENT-AD** | **A*T* contrast strategy** | | | | | |
|---|---|---|---|---|---|---|
| | **$A^{+}T^{*}$ & $A^{*}T^{+}$** | | **$A^{-}T^{-}$** | | **Statistic values** | |
| | n | % | n | % | χ2 | p |
| Sex | | | | | | |
| Male | 102 | 41 | 19 | 46 | 0.21 | 0.65 |
| Female | 146 | 59 | 22 | 54 | | |
| Race | | | | | | |
| Asian | 3 | 1 | 2 | 5 | 5.0 | 0.17 |
| Black | 7 | 3 | 3 | 8 | | |
| White | 232 | 94 | 35 | 85 | | |
| Unknown | 6 | 2 | 1 | 2 | | |
| | mean | std | mean | std | t | p |
| Age (year) | 67.8 | ± 7.7 | 70.4 | ± 6.5 | -2.1 | 0.04 |
| Education year | 15.6 | ± 2.8 | 16.5 | ± 2.4 | -2.1 | 0.04 |
